# Understanding accommodative control in the clinic: modeling latency and amplitude for uncorrected refractive error, presbyopia and cycloplegia

**DOI:** 10.1101/2023.03.20.533503

**Authors:** Jenny C. A. Read, Gerrit Maus, Clifton M. Schor

## Abstract

Accommodation refers to the process of increasing the optical power of the eye’s crystalline lens so as to focus on objects at different distances. Refractive error is a mismatch between the physical size of the eye and its optical power when accommodation is fully relaxed, while functional presbyopia is a decrease in the range of accommodation with age that results in the near point of accommodation lying beyond the near working distance. Both conditions mean that sharp focus may not be achievable for some distances, and observers will experience sustained defocus. A familiar example is an older person struggling to read text on their phone. Here, we identify a problem with current models of the neural control of accommodation: they predict excessive internal responses to stimuli outside the range of accommodation, leading to unrealistic adaptation effects. Specifically, after a prolonged period of viewing stimuli outside their accommodative range, current models predict that observers would show long latencies in the accommodative response to stimuli within range. These latencies are not observed empirically, indicating a problem with current models. We propose a simple solution, exploiting the predictive nature of accommodative control, and demonstrate that the new model performs correctly. We also model cycloplegia as a change in gain, and include a lower bound on the neural signal driving accommodation so as to model the additional relaxation of accommodation often seen with cycloplegia. We show that with these modifications, we can obtain plausible predictions for the accommodative response and accommodative convergence signal in a wide range of clinically relevant situations such as functional presbyopia, fogging lenses, corrected and uncorrected refractive error, and cycloplegia.

## Introduction

We recently published a model of neural control of accommodation, which aimed to bring together many ideas in the literature and to be reasonably comprehensive (Read et al., 2022). In particular, the model was the first to incorporate noise and to account for the closed-loop resonance enhancing responses at 1-2Hz.

We now wish to generalise the model to account for clinical situations such as functional presbyopia, refractive error, fogging lenses and cycloplegia. In previous models, e.g. that of Schor (1999) reproduced in Figure 1, the first two conditions have been modelled respectively by (a) adding a saturation nonlinearity after the plant transfer function, representing the limited range of accommodation, and (b) adding a constant offset, representing the refractive error (red elements in Figure 1). However, it has not previously been pointed out that this approach predicts a form of adaptation, in which viewing a stimulus beyond the accommodative range causes unrealistically long latencies in the accommodative responses to subsequent stimuli within the accommodative range.

**Figure 1.**
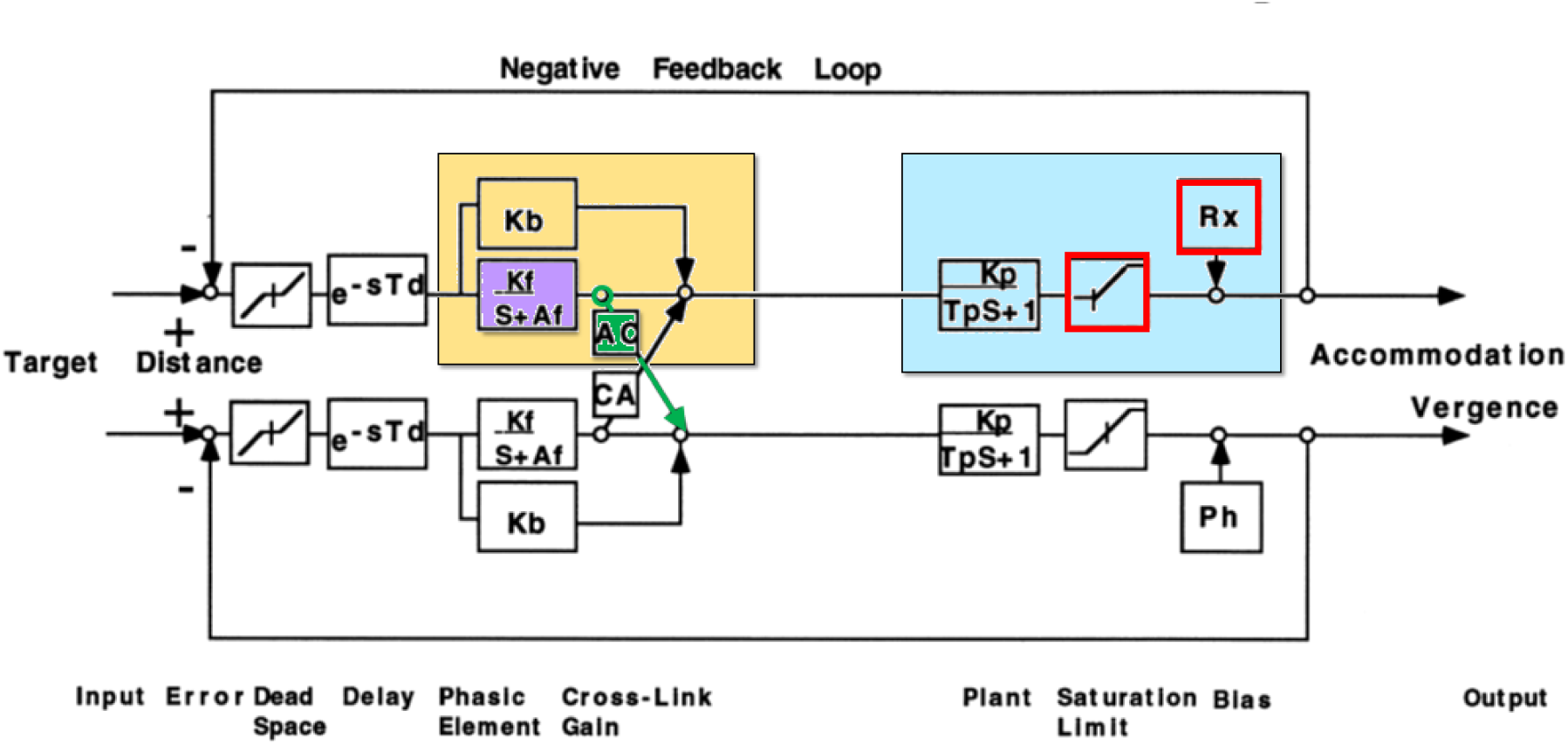
Reproduced from Figure 1b of Schor (1999), showing a model for the control of vergence and accommodation. To facilitate comparison with the present models, we have highlighted the accommodative control system in gold and the ocular plant in blue, as we do in diagrams of the present models. Note that since this model is non-predictive, the accommodative control system does not contain a virtual plant. We show this model to highlight how it handles the finite range of accommodation via a saturation block (red), the addition of refractive error (also red), and the accommodative-convergence AC signal (green) taken to be the output of the integrator (purple).

The basic reason for this behaviour is easy enough to understand: it stems from the very large internal signals predicted in response to sustained defocus. Consider for example the situation where an uncorrected +2D myope views a stimulus at infinity. The correct ocular power would be 0D, but the myope’s elongated eyeball means that their ocular power is +2D even when the lens is fully relaxed. They therefore experience a steady optical defocus of −2D (where optical defocus is defined as effective demand minus ocular power). Models of the neural control of accommodation are based on leaky-integral control, where the signal sent to the ocular lens is based on the integrated defocus encountered over the last few seconds. (In predictive models, the signal is actually based on the brain’s own estimate of defocus, but we can neglect this distinction for the moment.) In this example, the leaky integrator thus asymptotes to a large negative value, equal to the integrator’s gain multiplied by the steady-state defocus. The integrator’s gain must be large, around 8, in order to keep defocus errors small for stimuli within the accommodative range (Read et al., 2022), and so its asymptotic value for stimuli beyond range is large: −16D in this example. If the myope now transfers their gaze to a nearby object at +4D, the defocus will change from −2D to +2D. However, the signal sent to the lens will remain negative until the leaky integrator has fully discharged from its asymptotic value of −16D; only then will the lens begin to constrict as the signal begins to go positive. The response of accommodation to sinusoidal stimuli suggests a time-constant for the leaky integrator in the range 2.5s (Read et al., 2022). Thus, the latency before a response can be significant: well beyond the ∼300ms corresponding to the sensorimotor latency. In fact, when we simulate this example below (Figure 8i), we find that a delay of over 2s before any response begins.

While we have been unable to find published measurements of the latency of accommodative responses in uncorrected myopes, such long latencies are clearly unrealistic. They predict, for example, that a myopic child who has been trying to read the whiteboard several metres away at the front of the classroom would encounter a delay of some seconds before they could focus again on their own writing. Notice that the model is not simply predicting slower than usual responses: it is predicting a latency of some seconds before any response begins. This effect is also quite distinct from other forms of adaptation, such as Nearwork-Induced Transient Myopia (Hung & Ciuffreda, 2002, 1999). Current models make similarly unrealistic predictions for hyperopia and for functional presbyopia: in each case, exposure to a stimulus closer than the near point is predicted to cause large delays in the response to subsequent stimuli within range. Optometrists and people with refractive error can confirm that this is not their experience.

Current models also predict very long (seconds) latencies following defocus due to “fogging”. In clinical optometry, fogging is the practice of adding plus lenses in front of the patient’s eye so as to encourage accommodation to relax to the maximum amount possible (Ramsdale & Charman, 1988; Reese & Fry, 1941). This is particularly important in patients with hyperopia, who due to their refractive error will usually be accommodating even when viewing distant stimuli. The plus lenses effectively move the stimulus beyond the patient’s far point, making it appear blurred or “fogged”, and any accommodation makes this worse. Current models predict that when the plus lenses are removed and the person looks at a nearby object, there will be a delay of several seconds before the patient’s focus begins to return to normal. Again, this adaptation to fogging is not seen empirically.

Many people show a fixed minimum level of accommodation, which cannot be removed even with plus lenses. We know that this is due to a neural signal rather than the optical structure of the eye, since it can be substantially reduced when the ciliary muscle is paralyzed in pharmacological cycloplegia (Bagheri et al., 2018; Hiraoka et al., 2013; Yoo et al., 2017). This effect is again distinct from the adaptation predicted by current models.

The inability to simulate realistic responses following sustained defocus blur is a serious problem for current models of accommodation. It means they cannot correctly handle everyday clinical situations such as uncorrected refractive error, functional presbyopia and fogging lenses.

Although the focus of this paper is on accommodation alone, it is worth noting that this issue also presents some difficulty for current models of accommodative convergence. Empirically, when a normal observer views a stimulus with one eye, a change in stimulus distance from far to near elicits not only an accommodative response, but a nasalward movement of the occluded eye, so as to increase the convergence in a way that would help to null the retinal disparity if viewing were binocular. This accommodative convergence is taken to reflect a neural crosslink signal between the accommodative and vergence control systems. Many current models, e.g. the model of Schor (1999) shown in Figure 1, postulate that the accommodative-convergence or “AC” signal (shown green in Figure 1) is simply the output of the leaky integrator (purple) which controls accommodation. This normally works well. For example, if a functional presbyope is viewing a distant stimulus with one eye occluded, and an experimenter inserts a −1D lens, the accommodative integrator will increase its output by nearly 1D and both accommodation and vergence will increase accordingly. However, now suppose the same experiment is conducted closer than their near point, say with the stimulus at 4D. Accommodation will be unable to increase in response to the divergent lens, so the sustained defocus will charge the accommodative integrator to very high values. According to current models, therefore, vergence will thus increase by around 8 times more than it did previously for the same change in demand. Again, we simulate this example in detail in Figure 12 below, but the essential point is simple enough to be conveyed here. That is, current models predict a very large increase in the ratio of the accommodative convergence response/accommodation stimulus, i.e. stimulus AC/A ratio, for stimuli closer than the near-point. There is some empirical support for such an effect (Alpern et al., 1959), but not really of the predicted magnitude. Larger increases in stimulus AC/A ratio are, however, seen with partial and complete cycloplegia. In partial cycloplegia, the accommodative amplitude gradually decreases while the AC/A ratio gradually increases as the depth of cycloplegia increases (Christoferson & Ogle, 1956), and in complete cycloplegia, the amplitude of accommodation reaches zero and the AC/A becomes nearly infinite (Morgan, 1954).

In this paper, we present an adjustment to previously published models which addresses all these problems. Our model can produce more realistic simulations of accommodation in the presence of refractive error, fogging lenses and functional presbyopia, while keeping the output of the accommodative integrator more consistent with empirical constraints on the accommodative-convergence signal. We also model the relaxation of accommodation seen with pharmacological cycloplegia, where the ciliary muscle is relaxed more completely than is otherwise possible and so the plant assumes the minimum possible optical power.

As in our previous paper, the key to achieving this is predictive control: the system’s ability to model and predict its own responses. We postulate that current models are incorrect in assuming that all defocus error inevitably charges up the leaky integrator controlling accommodation, as shown in Figure 1. Rather, we postulate that as part of predictive control, the brain is able to recognise when defocus occurs because the stimulus is further away than the accommodative far point, and avoids feeding such uncorrectable defocus into the integrator which generates the neural control signal. For stimuli closer than the maximum ocular power, we postulate that the resulting defocus is initially fed through into the integrator, generating a neural control signal asking for more accommodation than is physically possible for the plant. However, we suggest that the brain internally reduces its estimate of correctable defocus based on the neural control signal rather than the physical accommodation. The result is that, again, the steady-state integrator output and neural control signal remain fairly small even after sustained defocus.

Adopting different approaches for defocus due to stimuli that are too far versus too close enables us to keep the output of the integrator consistent with accommodative convergence. In our model, when a monocular stimulus beyond the observer’s far point moves still further away, no accommodative divergence movement is triggered. But when a monocular stimulus closer than the near point comes closer still, an accommodative convergence movement is triggered.

For ease of reference, our final proposed model is shown here in Figure 2. In the Methods, we go through the different components and explain the reasoning behind the modifications. In the Results, we show simulations side-by-side for the original model and for our proposed new model, demonstrating both the problems with the original model and how our new model fixes them.

**Figure 2.**
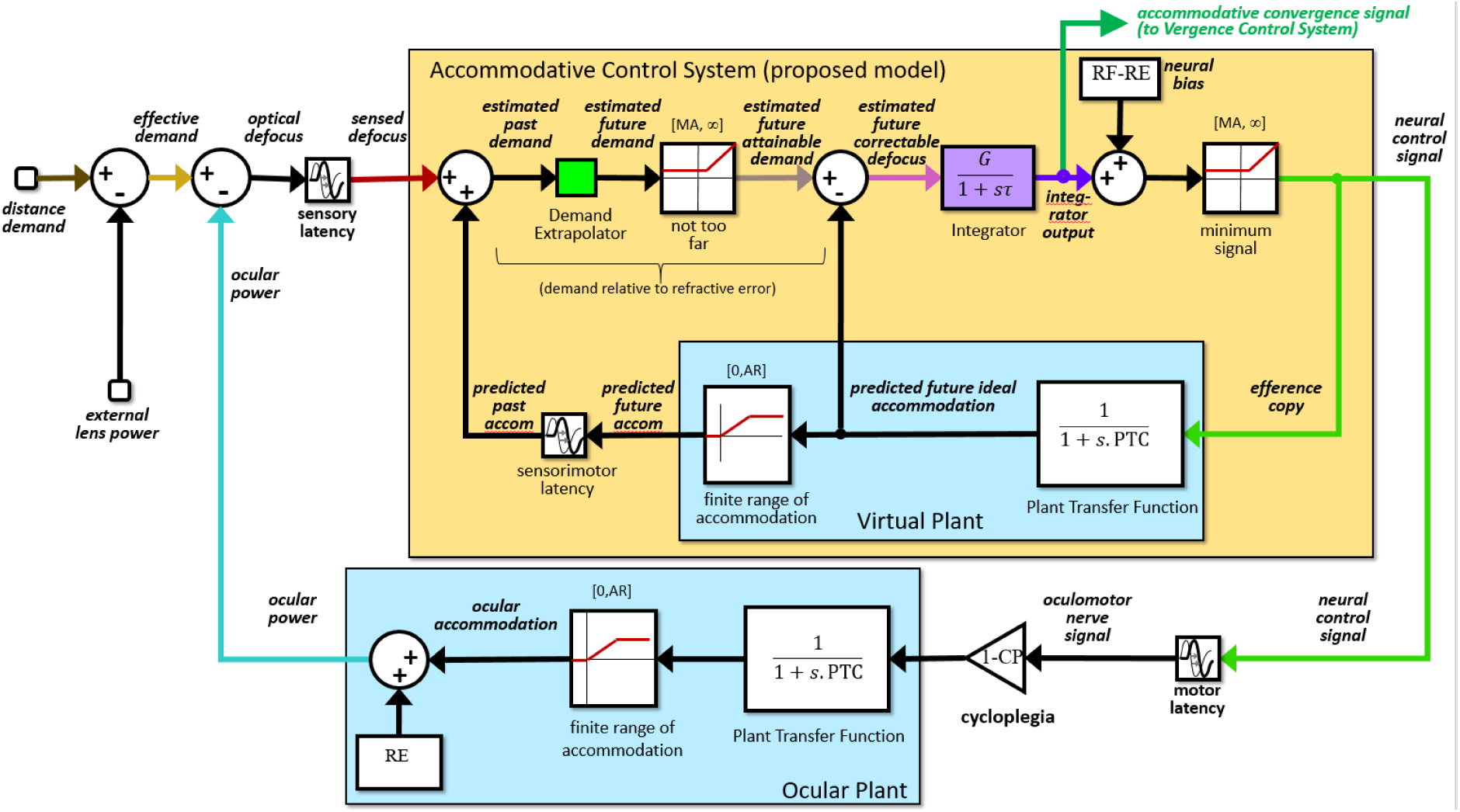
Diagram of the full proposed model, showing how external lenses and cycloplegia are accounted for. The colors used for the distance demand, effective demand, ocular power, sensed defocus, estimated future attainable demand, estimated future correctable defocus, integrator output and neural control signal match the colors used to plot these signals in the results figures. The green arrow at the top shows where the accommodative convergence (AC) signal is drawn off and fed into the Vergence Control System (not modeled in this paper).

## Methods

All the models in this paper, as well as the full model in Read et al (2022), conform to the general structure shown in Figure 3. This describes a negative feedback loop in which the block labelled “Accommodative Control System” generates a control signal which is sent to the block labelled Ocular Plant (the ciliary muscle, lens etc) so as to generate an accommodative response which aims to match stimulus demand and thus minimise defocus error. This is reviewed in detail in Read et al (2022). Note that while in the earlier paper the output of the plant was described as “accommodation”, we now refer to it as “ocular power”. This is because, in the presence of refractive error, ocular power is not in general equal to accommodation (e.g. the optical power of a myopic eye is >0D even when accommodation is fully relaxed with complete cycloplegia).

**Figure 3.**
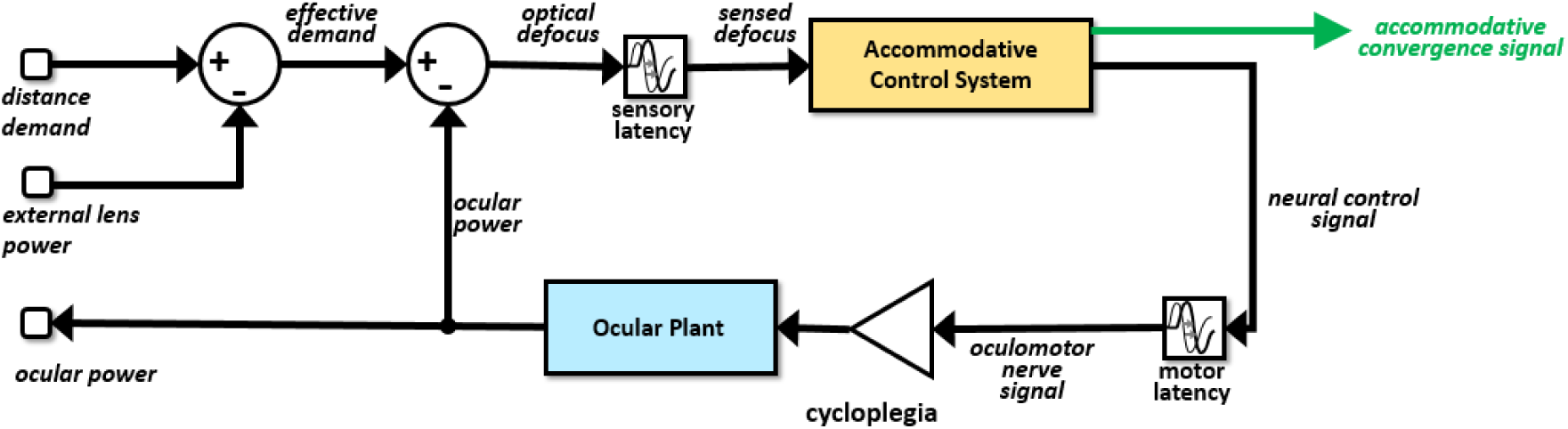
Overall model structure. Stimulus distance demand and any external lenses (both in diopters) combine to produce an effective stimulus demand. The difference between this and ocular power, i.e. the optical power of the eye, gives the optical defocus. This in turn is sensed by the brain following a sensory latency allowing time for defocus to be computed. The accommodative control system takes the sensed defocus as input and outputs a neural control signal for accommodation. For completeness we also show the accommodative convergence signal (green) which goes into the vergence control system, not considered in detail in this paper. The neural control signal for accommodation travels down the oculomotor nerve and arrives at the ocular plant after the motor latency. We label the signal “neural control signal” when it is computed in the brain, and “oculomotor nerve signal” when it arrives at the plant. Cycloplegia e.g. with atropine or cyclopentolate effectively inhibits this signal, as indicated by the triangular gain block. In complete cycloplegia, gain is reduced to zero and effectively removes all input to the plant (i.e. cuts the signal). In partial cycloplegia, gain is reduced and large steady state errors manifest as a reduced amplitude and elevated AC/A ratio. The ocular power is the output of the ocular plant. The internal structure of the blue Ocular Plant block is shown in Figure 4. Possible internal structures for the yellow Accommodative Control System block are discussed in Figure 5, Figure 15 and Figure 7. Figure 7 shows the structure we are proposing in this paper.

The internal structure of the Ocular Plant block is shown in Figure 4. It receives as input the signal to the ciliary muscle from the oculomotor nerve, and gives as output the optical power of the eye. The basic transfer function is a leaky integrator, equivalently a first-order low-pass filter, specified by its time-constant, the model parameter *PlantTimeConstant*, abbreviated PTC in the figures, which in this model is set to 156ms (Table 1). The saturation block after the plant transfer function imposes the finite range of accommodation. A saturation block is a non-linear circuit element which has lower and upper limits [*L,U*]. Inputs within this range are passed through unchanged; for inputs less than its lower limit *L* or greater than the upper limit *U*, the output is *L* or *U* respectively. In this case, the lower limit is 0D and the upper limit is the model parameter *AccomRange*, abbreviated AR, also in diopters D. This represents the finite range of accommodation that declines with age. Finally, we add on *RefractiveError* or RE, the observer’s refractive error, also in diopters D. Note that in this paper, we are considering the refractive error of the eye, rather than the power of the lens used to correct it. Ocular refractive errors are therefore *positive* for myopes (a myopic eye has too much optical power for its length and needs correction with negative lenses) and *negative* for hyperopes.

**Figure 4.**
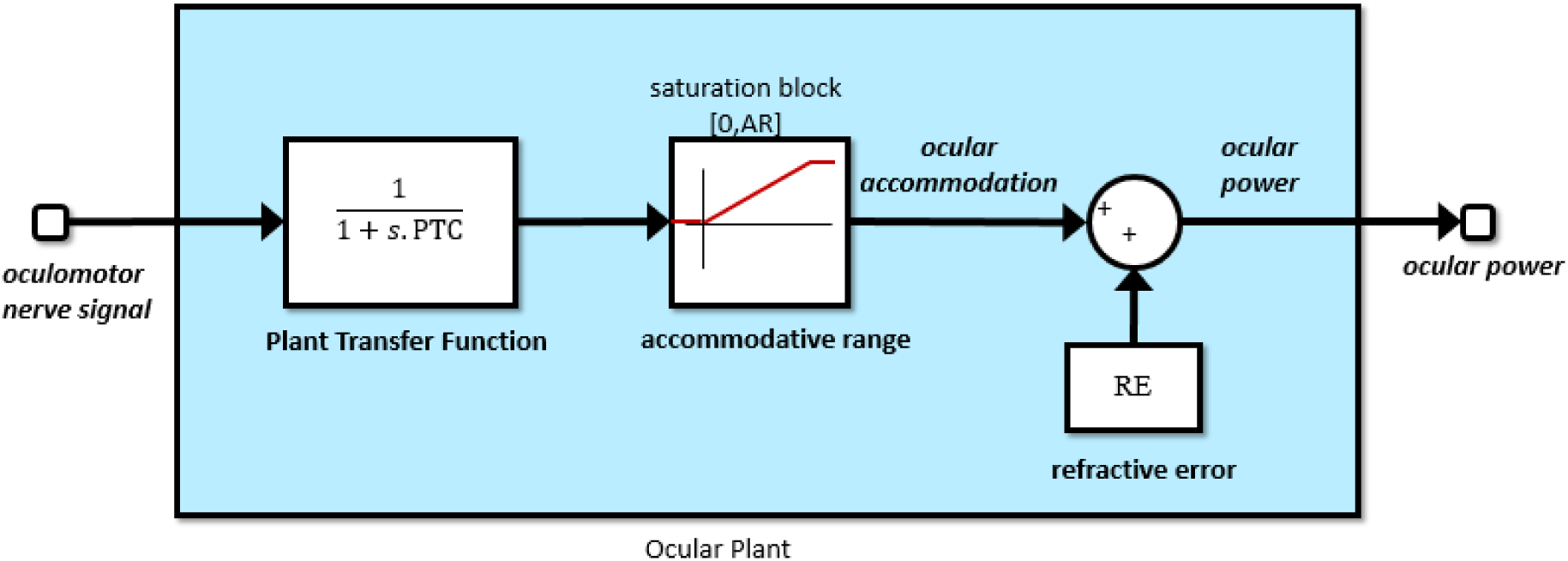
Structure of the Ocular Plant block shown in Figure 3. The basic transfer function is a leaky integrator or first-order low-pass temporal filter, with a time-constant given by the model parameter PlantTimeConstant (Table 1). This is followed by a saturation block representing the finite accommodative range, set by model parameter AccomRange. Inputs in the range [0,AccomRange] are passed unchanged; inputs less than zero or greater than AccomRange produce outputs of zero or AccomRange respectively. Finally, the model parameter RefractiveError is added. PTC=PlantTimeConstant, AR=AccomRange, RE=RefractiveError (Table 1).

**Table 1.** Parameter values for the Simulink model supplied with the paper and used to obtain the results (except where stated otherwise). These values are visible in the Simulink Model Workspace, and can be altered there if desired. Notice that the actual rest focus of the model, i.e. the asymptotic ocular power in open-loop mode, is equal to the parameter RestFocus only when RestFocus>(MinAccomSignal+RefractiveError). When the parameter RestFocus is less than this, the model’s actual rest focus is equal to (MinAccomSignal+RefractiveError).

| Parameter | Name in Simulink workspace | Abbreviation in figures | Value |
| --- | --- | --- | --- |
| Rest focus | *RestFocus* | RF | 1.4D |
| Sensory latency | *SensoryLatency* |  | 0.20s |
| Motor latency | *MotorLatency* |  | 0.10s |
| Time constant of plant | *PlantTimeConstant* | PTC | 0.156s |
| Gain of integrator | *FastGain* | G | 8.0 |
| Time constant of integrator | *FastTimeConstant* | $\tau$ | 2.5s |
| Minimum value of the neural control signal for accommodation | *MinAccomSignal* | MA | 0.2D |
| Refractive error (as measured under cycloplegia) | *RefractiveError* | RE | 0D or as stated |
| Accommodative range | *AccomRange* | AR | 4D or as stated |
| Cycloplegia | *Cycloplegia* | CP | 0 or as stated |

As an example, consider an observer with *AccomRange*=4D, typical for someone in their late forties. An emmetropic observer with this range can focus on stimuli from 0D to 4D (infinity down to 25cm). A myope with +2D of refractive error can focus on stimuli from 2D to 6D (50cm to 17cm), while a −2D hyperope can focus on stimuli from −2D to 2D (i.e. can focus unaided on stimuli from infinity down to 50cm, and can also tolerate plus lenses of up to +2D without blur). All model parameters are given for reference in Table 1.

### Simplified model of accommodative control used in this paper

For simplicity, in this paper we will use a stripped-down version of the model developed in Read et al (2022), removing elements that are irrelevant to the point at issue. To this end, we remove noise, the clipped proportional signal and the slow integrator, as well as continuing to neglect the pulse component of the response to step changes. We retain a bias signal controlling rest focus. (If we retained the slow integrator, the latencies would potentially be even longer, since both the slow and the fast integrator would charge up.)

Figure 5 shows the contents of the “Accommodative Control System” block in Figure 3 as it would be in this stripped-down model. This contains a virtual internal model of the accommodative plant, shaded light blue in the figures, which is used to generate internal predictions about future ocular power, demand and defocus. See our earlier paper for an in-depth discussion of this predictive control. In brief, the input signal to the accommodative control system is the sensed defocus, assumed to be computed from retinal information such as blur and higher order aberrations, longitudinal chromatic aberrations etc. However, to avoid control problems such as overshoot and ringing, the neural control signal is not generated directly from this sensed current defocus, but from the brain’s estimate of the likely *future* defocus, taking into account the predicted changes in accommodation based on the control signal already sent to the eye. These are computed via an efference copy of the neural control signal, which is fed into a virtual plant model, outlined in light blue. The output of the virtual plant is the eye’s predicted future optical power. Figure 5 shows how this is delayed and then added onto the sensed defocus to compute the predicted past demand. The green block labeled “Demand Extrapolator” in Figure 5 uses this to estimate the future demand. This block was referred to as the “Demand Predictor” in our previous paper. We now feel the term “Demand Extrapolator” is more helpful, as it emphasizes that stimulus demand is not necessarily within the control of the observer and thus cannot in principle always be perfectly predicted. In our Simulink models, the Demand Extrapolator in fact simply feeds through its input, i.e. it assumes the effective demand will remain constant for at least a time equal to the sensorimotor latency. The predicted future power is then subtracted from this estimated future demand to result in the estimated future defocus, which is the key signal used for accommodative control.

**Figure 5.**
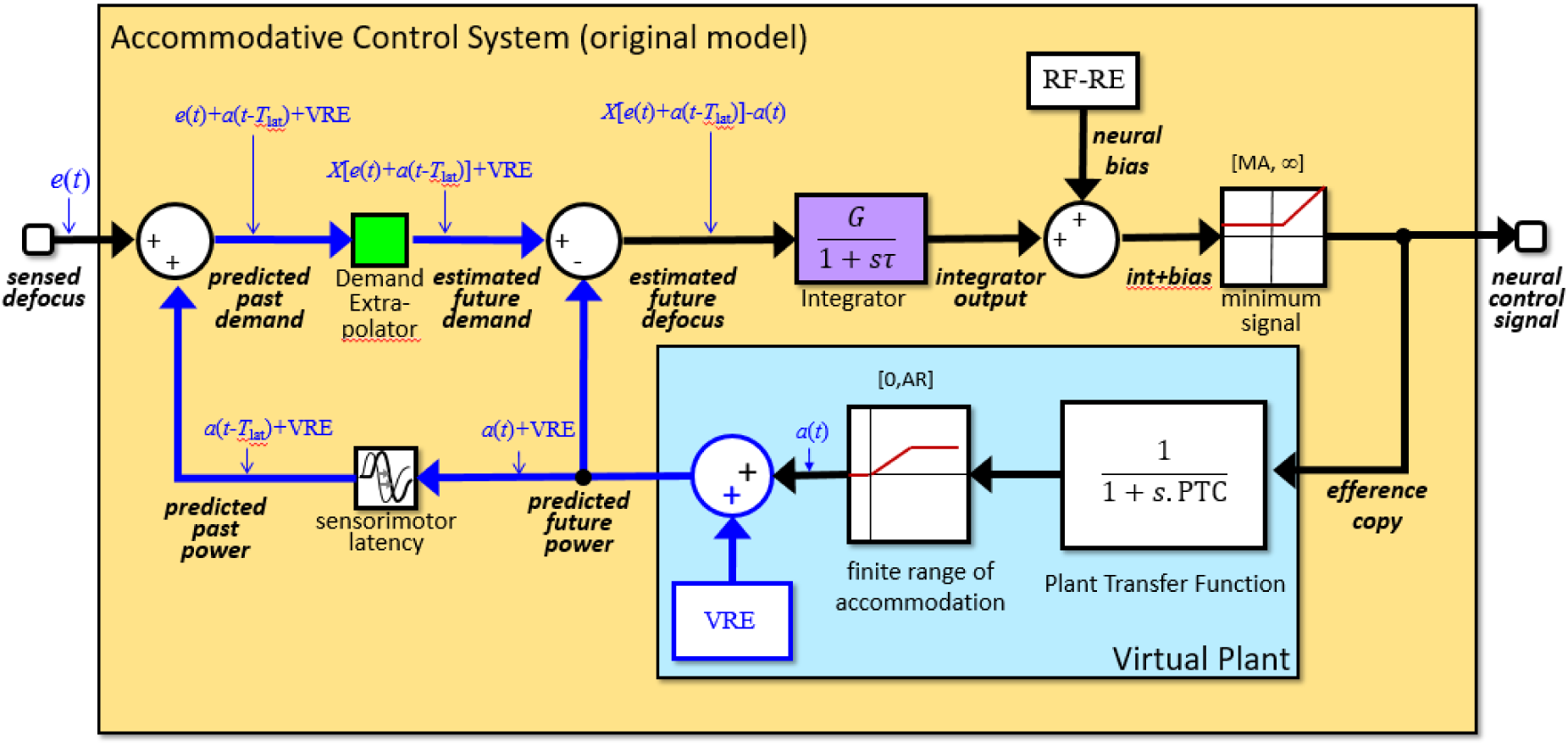
Structure of the Accommodative Control System (yellow block in Figure 3) in the version we refer to as the “original model”. This is as in Read, et al (2022), but with refractive error and a minimum neural accommodative signal added and with extraneous elements (noise, slow integrator, proportional signal) not relevant to this paper removed for simplicity. Signals drawn in blue are those affected by the “virtual refractive error”, VRE, inside the blue Virtual Plant block. Apart from this virtual refractive error, the virtual plant is identical to the Ocular Plant block shown in Figure 4. The output of the green Demand Extrapolator block is shown as a general function X, which is assumed to be linear so that X(input+constant)=X(input)+constant. In our Simulink models, X is the identity, i.e. the Demand Extrapolator simply assumes that the stimulus demand will remain at its current value. Here and subsequently, we use the adjective “predicted” (e.g. predicted future power) to refer to the control system’s computation of external signals which are error-free in our model (since they depend only on the system’s own behavior and on the properties of the ocular plant, which could in principle be learnt by the brain and are known to our model). We use the term “estimated” (e.g. estimated future demand) to refer to the control system’s computation of external signals which are not fully within its control, e.g. because they depend on the self-motion of scene objects. RF,RE are abbreviations for the model parameters RestFocus, RefractiveError. VRE = “virtual refractive error”. As discussed in the text, where demand extrapolation is linear, VRE can be set to zero without loss of generality. Notice that here and subsequently, to limit clutter we do not show the accommodative convergence signal, but it remains equal to the integrator output.

The estimated future defocus is fed into the Integrator block, shaded purple in the figures. This corresponds to the “fast” integrator in the full model. It is a leaky integrator which is the heart of the model’s control of accommodation. Its gain and time-constant are central to the model’s behavior.

The model also includes a neural bias signal, which is added onto the integrator output. As discussed below, this bias helps set the rest focus adopted in open-loop viewing.

In the Results section we will demonstrate that the original Accommodative Control System shown in Figure 5 predicts unrealistic latencies, and we will discuss various alternatives. Before proceeding to this, it will be helpful to discuss various aspects which apply to all models discussed in the paper.

### Modeling the relaxation of accommodation with cycloplegia

In Figure 5, the sum of the bias and integrator signals passes through a saturation block, marked “minimum signal”, before the resulting neural control signal is passed to the oculomotor nerve and to an internal efference copy. This saturation block has no upper limit (i.e. the upper limit is set to infinity). In our previous paper (Read et al., 2022), its lower limit was set to zero, ensuring that only positive signals were sent to the ciliary muscle. This models the fact that once the ciliary muscle is fully relaxed, no further relaxation can occur, or equivalently that accommodation cannot drop below zero.

For this paper, we introduce a new parameter, *MinAccomSignal*, abbreviated MA (Table 1). This non-negative parameter forms the lower limit of the “minimum” saturation block. It represents the non-zero level of accommodation observed in most people, even emmetropes, when they are viewing stimuli at infinity, or effectively beyond infinity by the use of plus lenses. We know that this must result from a neural signal because when the ciliary muscle is paralyzed pharmacologically, effectively cutting the oculomotor nerve signal, accommodation typically drops below the value obtained with stimuli at or beyond infinity (Bagheri et al., 2018; Hiraoka et al., 2013; Yoo et al., 2017). Including this additional *MinAccomSignal* parameter enables us to model this well-known effect. If the input to the saturation block exceeds *MinAccomSignal,* the output is just the input; if it is less than *MinAccomSignal*, the output is set equal to *MinAccomSignal*.

### Modeling rest focus via a neural bias

In open-loop mode with the defocus signal clamped at zero, e.g. when viewing through pinholes, accommodation does not settle to zero but to a finite value. This implies a signal that operates even in the absence of any input to the system. Our *MinAccomSignal* parameter, discussed above, could account for this, but would imply that the rest focus would be equal to the accommodation measured when viewing a distant stimulus, plus any refractive error. In fact, rest focus is usually much larger, around 1.4D, even after correction for refractive error (Fisher et al., 1987; McBrien & Millodot, 1987; Rosenfield et al., 1993). To account for this, we include a constant neural bias signal, which is added on to the output of the integrator when generating the neural control signal.

In models with no refractive error, the rest focus is equal to the neural bias (provided that the bias is greater than the minimum neural signal *MinAccomSignal*). In models with refractive error, the refractive error itself effectively acts as an external bias. Thus, the rest focus is equal to the neural bias plus the refractive error (again, provided that this is greater than the minimum neural signal).

We could include a model parameter representing neural bias. However, for consistency with our previous paper, we instead parametrise this via a model parameter *RestFocus*, and set the neural bias to *RestFocus-RefractiveError*, as shown in Figure 5. If the model parameters satisfy *RestFocus>RefractiveError+MinAccomSignal*, then open-loop ocular power will asymptote to the parameter *RestFocus.* If this condition is not met, open-loop ocular power will asymptote to *RefractiveError+MinAccomSignal*, i.e. the minimum ocular power possible without cycloplegia.

### Note on the definition of tonic accommodation

“Tonic accommodation” is often used to refer to the neural signal driving the rest focus measured in darkness or in open loop (McBrien & Millodot, 1987; Rosenfield et al., 1993). This would correspond to our neural bias term, *RestFocus-RefractiveError*. However, other workers define “tonic accommodation” as “the eye’s normal functional state for distance” (Hiraoka et al., 2014), which in our model is represented by *MinAccomSignal*. Since there is no single “tonic accommodation” in our model, we avoid using the term.

### Note on the definition of refractive error

In this paper, we define “refractive error” as the optical power of the eye when accommodation is fully relaxed. However, our *MinAccomSignal* parameter means that accommodation cannot be fully relaxed unless the eye is completely cyclopleged. It follows that our *RefractiveError* parameter represents the optical power of the eye under cycloplegia. In our model, the optical power measured when the eye views an object at infinity, or through plus lenses, is *RefractiveError+MinAccomSignal*. In some empirical studies, what is referred to as “refractive error” may therefore correspond in our model to refractive error plus this minimum neural signal.

### The brain does not need to know its own refractive error

In Figure 5, the virtual plant model is shown as including a virtual refractive error term, VRE. However, the value of this term does not affect the behavior of the model, at least when the Demand Extrapolator block is linear. To show this, the small blue labels in Figure 5 mark on the values of selected signals, starting with the accommodation output by the virtual plant, *a*(*t*), and the sensed defocus error, *e*(*t*). As shown in Figure 5, the value of VRE affects the predicted ocular power and thus the estimate of demand fed into the Demand Extrapolator block. Let’s denote the function performed by the Demand Extrapolator block as X(*input*). If the function *X* is linear, then by definition X(*input*+VRE)=X(*input*)+VRE. The VRE term then cancels out when the predicted future power is subtracted from the estimated future demand. Thus, the estimated future defocus, drawn pink in Figure 5, is independent of the value of VRE. The value of VRE affects the internal signals shown in blue, but has no effect on the overall transfer function of the Accommodative Control System. VRE can thus without loss of generality be set to zero, and we will adopt this simplification in the rest of the paper.

Similarly, we saw above that it was convenient to write the neural bias signal as *RestFocus*-*RefractiveError.* This again does not mean that the brain necessarily “knows” the eye’s refractive error. The brain simply applies a neural bias, and the refractive error adds on to this to determine the system’s rest focus.

Thus, none of the models discussed in this paper require us to postulate that the brain has any information about the ocular refractive error.

### Developing a new model

As we show in the Results section, the problems with current models stem from the same basic problem: excessive integrator output during exposure to stimuli beyond the accommodative range, which then leads to long latencies to subsequent stimuli within range. To fix the issues, we assume that neural control is sophisticated enough to recognise that sustained charging in such a situation is unhelpful, and takes steps to prevent it occurring. Defocus which cannot be corrected, due to ocular limitations, is ignored and not used to charge the integrator; only *correctable defocus* is passed on to the integrator.

#### Driving accommodation with estimated future correctable defocus

This can be achieved by changing the Accommodative Control System from the model shown in Figure 5 to that shown in Figure 6. The key difference from the original model is the additional saturation block, labeled “finite range” and outlined in red in Figure 6. This converts estimated future demand into estimated future *attainable* demand. The lower limit of this saturation block is the *MinAccomSignal* parameter, while its upper limit is *AccomRange*. When demand is within range, therefore, the model operates exactly as before. But when demand is out of range of the observer, the saturation block prevents uncorrectable defocus error from charging the integrator up to its asymptotic value.

**Figure 6.**
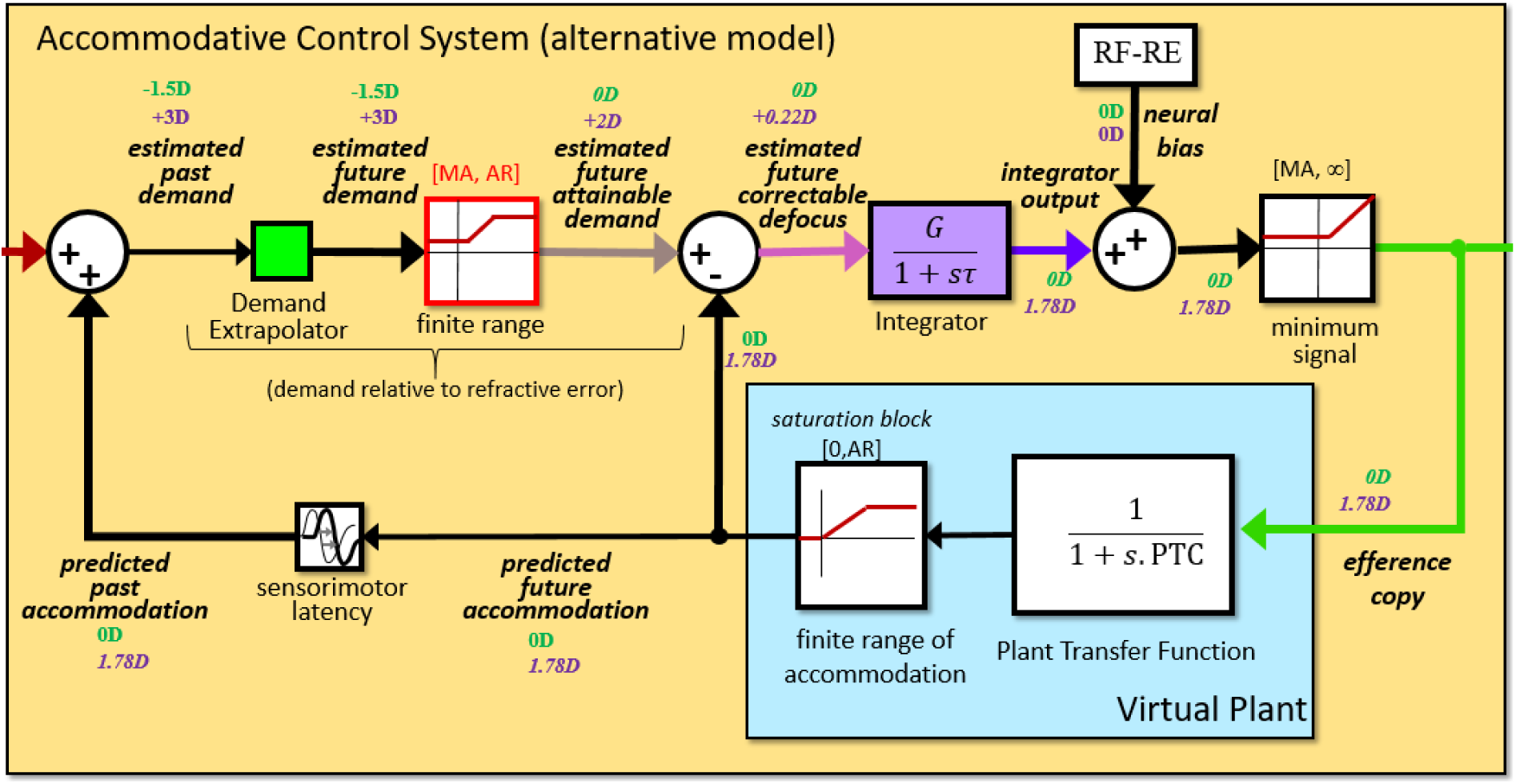
As Figure 5, but we have added a saturation block (red) clipping estimated future demand at values attainable by the plant. This means that the signal passed to the integrator is the correctable component of the estimated future defocus. We have also removed the refractive error term from the virtual plant, since as shown above this does not change behavior. In consequence, the signals labeled “demand” actually represent “demand minus refractive error”. This model solves the problem of long latencies but at the cost of removing the accommodative convergence signal for stimuli out of accommodative range. Signals shown in color match colors used in results figures.

For example, consider an older emmetrope, with a maximum ocular power of +2D, viewing a stimulus at +3D. The model of Figure 6 predicts steady-state accommodation of 1.78D (i.e. 8/9 of 2D, where the 8 is the closed-loop gain of the system). This results in steady-state defocus of 1.22D. In this new model, the estimated attainable future demand is clipped to +2D. The estimated future *correctable* defocus is therefore 2D-1.78D = 0.22D. This is the steady-state input to the integrator which, when multiplied by the gain, produces the steady-state accommodation of 1.78D. In contrast, in the original model shown in Figure 5, steady-state defocus of 1.22D would have charged the integrator up to 8*1.22D or nearly 10D.

#### Accommodative convergence in response to stimuli out of accommodative range

The model of Figure 6 therefore solves the problem of excessive steady-state integrator output leading to excessive latencies. However, it causes another problem. Let’s go back to the previous example of the older emmetrope, and consider what happens if the stimulus moves still closer, say from +3D to +4D. The answer is that nothing happens to the integrator or the motor signal. The sensed defocus will of course increase by 1D, from 1.22D to 2.22D, so the estimated future demand will increase to 4D. But the estimated future *attainable* demand will still be 2D, and so none of the other signals in the system will change.

This raises a question about the accommodative-vergence crosslink. Efforts of accommodation that exceed the amplitude of accommodation still stimulate accommodative convergence. In addition, convergence continues to increase with efforts of accommodation after reaching the full amplitude of accommodation *and* disparity convergence, presumably due to efforts of accommodation and accommodative convergence (van der Hoeve & Flieringa, 1924). Thus, the increase in defocus due to the approach of a monocular stimulus, even beyond the near-point, causes a detectable vergence response even though accommodation cannot change. In this paper, we are not including the full accommodation-vergence model, but we still want to be aware of where the accommodative-convergence signal might come from. In previous models, the accommodative-convergence signal is the output of the accommodative integrator, as in Figure 1. However, the output of the integrator in Figure 6 is not suitable for this purpose, since it predicts no accommodative-convergence signal when stimuli move closer than the accommodative near point.

One approach would be to postulate a different integrator for accommodative convergence, but that seems an unnecessary complication. Instead, we choose to adopt the simpler approach shown in Figure 7.

**Figure 7.**
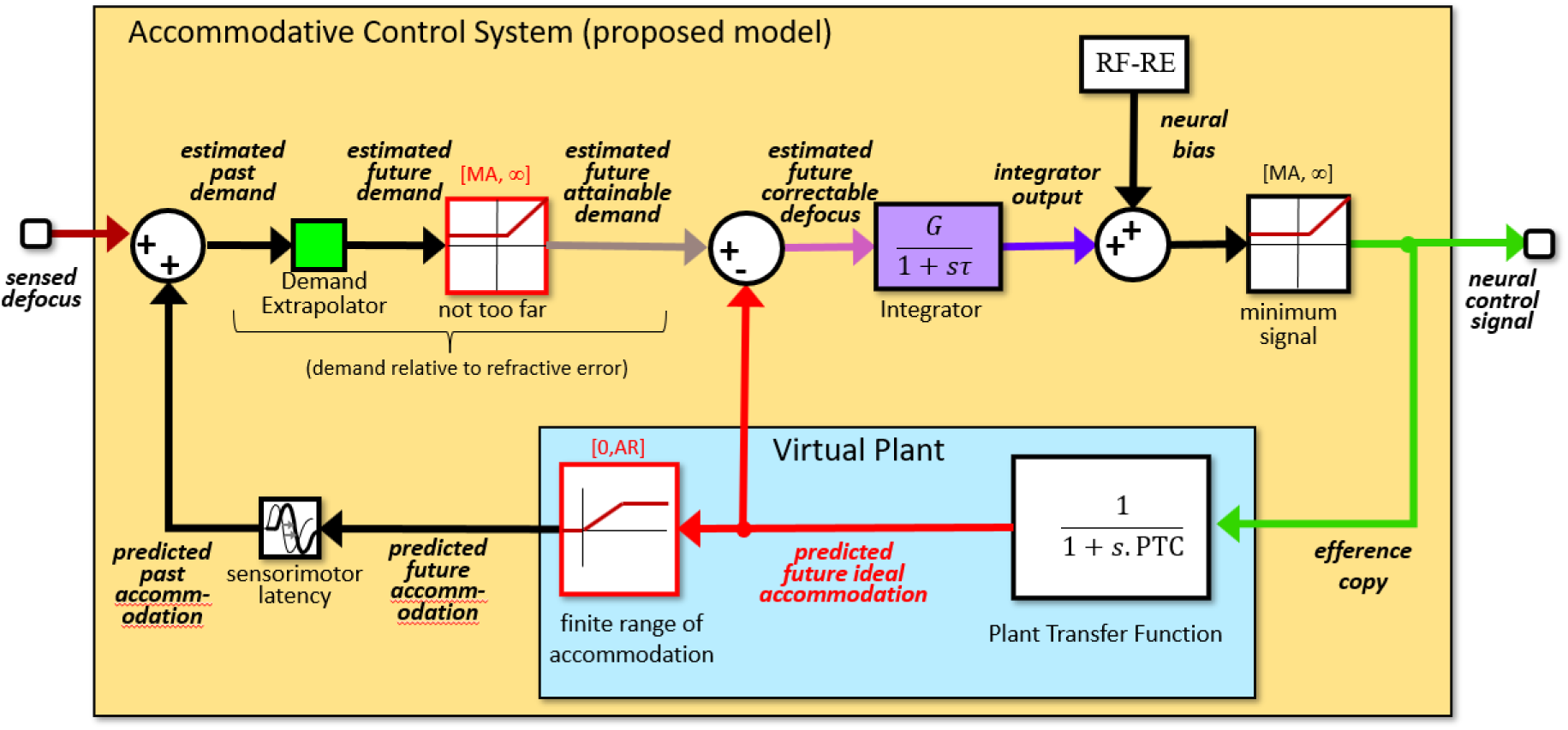
Accommodative Control System of the proposed new model. As Figure 6 but the “finite range” saturation block now has no upper bound, and is now labeled “not too far”. Additionally, there are effectively now two virtual plants: one outputting the predicted future accommodation, with the same finite range as the physical plant, and one outputting the ideal accommodation, with no upper limit.

#### Our proposed model

Figure 7 makes two changes relative to Figure 6, all highlighted in red; for reference, Figure 2 presented the full model with this Accommodation Control System. First, the saturation block after the Demand Extrapolator now accounts for only the lower bound on accommodation. The attainable demand is still lower-bounded by *MinAccomSignal*, but is now not upper-bounded. The effect of this is that demand that is too distant to focus on is clipped, as in Figure 6, but demand from stimuli that are too close is fed through into the defocus computation, as in the original model. To prevent such unattainable demand from driving the integrator to excessive values, when computing the future correctable defocus, we now subtract off the “predicted future *ideal* accommodation” signal, which also ignores the observer’s near-point. Note that in this new model the brain does still compute the actually-attainable predicted future accommodation, taking into account the finite accommodative range, and this is used as before to reconstruct the demand relative to refractive error. However, now a “predicted future ideal accommodation signal” is read out directly after the virtual plant transfer function, before the saturation block labeled “finite range of accommodation”. This “ideal” signal is still lower-bounded by *MinAccomSignal*, since the efference copy input to the plant is also lower-bounded there, as indicated by the saturation block labeled “minimum signal”.

The modifications made between Figures 6 and 7 make no difference to the behavior for stimuli that are too distant for the observer to focus on. However, they do affect the response to stimuli that are too close. First, the removal of the upper bound on the saturation block computing the estimated future attainable demand means that the accommodative control system continues to track changes in stimuli that are too close, i.e. where demand is already higher than the maximum ocular power.

Second, the construction of an ideal accommodation signal which also has no maximum power ensures that the initial transient response to such changes can be correctly integrated and then nulled. Together, this ensures that the change in integrator output is roughly equal to the change in demand, not many times greater as in the original model. Subsequent latencies therefore remain close to normal, but the integrator output can still be used to drive accommodative vergence. Finally, the measurable ocular power or accommodation response is limited to the near point by the upper bound or limit of the saturation block labeled “finite range of accommodation” that appears in both the internal and external feedback loops shown in Figure 2.

### Simulation details and model parameters

Simulations were run in Matlab Simulink, R2019a, using a fixed-step solver with timestep set to 1ms. All models and code are provided with the paper, and are available to download at https://figshare.com/s/99f9653e56d02136272c

## Results

### Current models produce unrealistically long latencies following stimulation beyond the accommodative range

The central topic of this paper is the unrealistically long latencies observed in current models after exposure to demands beyond the accommodative range of the simulated observer. This is demonstrated in Figure 8. Here and in subsequent figures, results for the original model are shown on the left while results for our proposed new model are shown on the right, to facilitate comparison.

**Figure 8.**
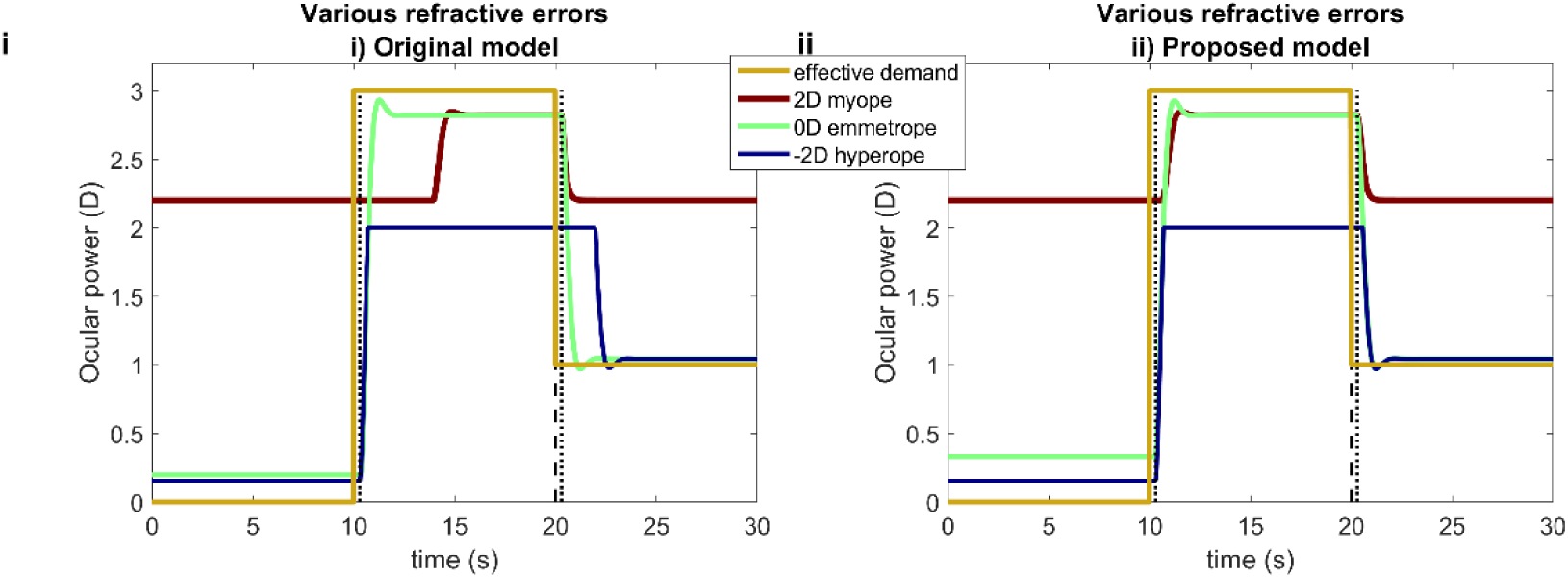
Time-courses of ocular power for a stimulus which is initially at infinity (0D), then moves to 33cm (3D) at t=10s, then moves back to infinity at t=20s, for three different functionally presbyopic observers with accommodative range 4D and refractive errors indicated in the legend, for (i) the original model, Figure 5, and (ii) our proposed new version, Figures 2 and 7. Dashed lines mark the time of step changes in demand, and dotted lines these plus the sensorimotor latency of 300ms. This figure was obtained with Matlab file PlotOcularPowerDifferentRE.m

In Figure 8, the time-course of the stimulus effective demand is shown in yellow. It steps from 0D at time zero up to +3D at t=10s and back down to 1D at t=20s. The accommodative response is shown for three simulated observers with different amounts of refractive error: an emmetrope with 0D, a myope with +2D, and a hyperope with −2D. All simulated observers are assumed to have the same total accommodative range, 4D, and the same minimum neural accommodative signal, 0.2D.

The emmetrope and hyperope accommodate close to 0D for the 0D stimulus, but the myope only gets to around +2.2D, reflecting their refractive error plus minimum neural accommodative signal. They thus experience defocus while viewing this distant stimulus. When the stimulus demand steps up to 3D at t=10s, the emmetrope and myope both successfully get close to 3D, but for the −2D hyperope, +3D is out of range. Finally, when the stimulus demand steps down to 1D at t=20s, the emmetrope and hyperope successfully accommodate close to this value, but the +2D myope cannot focus on objects this far away.

These steady-state values are as expected for the specified refractive errors, accommodative range, gain and minimum neural accommodative signal. However, notice the latencies for the original model (Figure 8i). At t=10s, the emmetrope and hyperope begin responding immediately after the expected sensorimotor latency of 300ms. However, the myope does not begin responding until some seconds after the step change in demand. Similarly when the demand drops to 1D at t=20s, while the other simulated observers begin relaxing accommodation after 300ms as expected, the hyperope does not begin responding until 1700ms after the stimulus change. As discussed, these long latencies are extremely unrealistic and thus indicate a problem with current models. Figure 8ii confirms that our proposed new model does not show these long latencies.

### The excessive latencies are due to excessive integrator output in current models

Figure 9iA illustrates the underlying reason for the long latencies in the original model, for the example of an uncorrected +2D myope observing a stimulus which is initially at a distance of 4D, then at t=5s switches to 0D, then at t=20s switches back to 4D. This demand time-course is shown in Figure 9iA by the thick brown line with yellow dots. The remaining curves show various signals for the myopic observer. The heavy blue line shows the time-course of the ocular power, and the heavy green line shows the time course of the motor signal which is 2D lower than the ocular power due to the myopic refractive error. When the stimulus is at 4D, the ocular power is close to 4D (a little less due to accommodative lag), so the stimulus is in focus. When the stimulus moves to 0D at t=5s, the myope relaxes accommodation, but due to their +2D refractive error they cannot bring the ocular power below +2D. They therefore experience a constant defocus error of −2D (red, pink traces in Figure 9iB).

**Figure 9.**
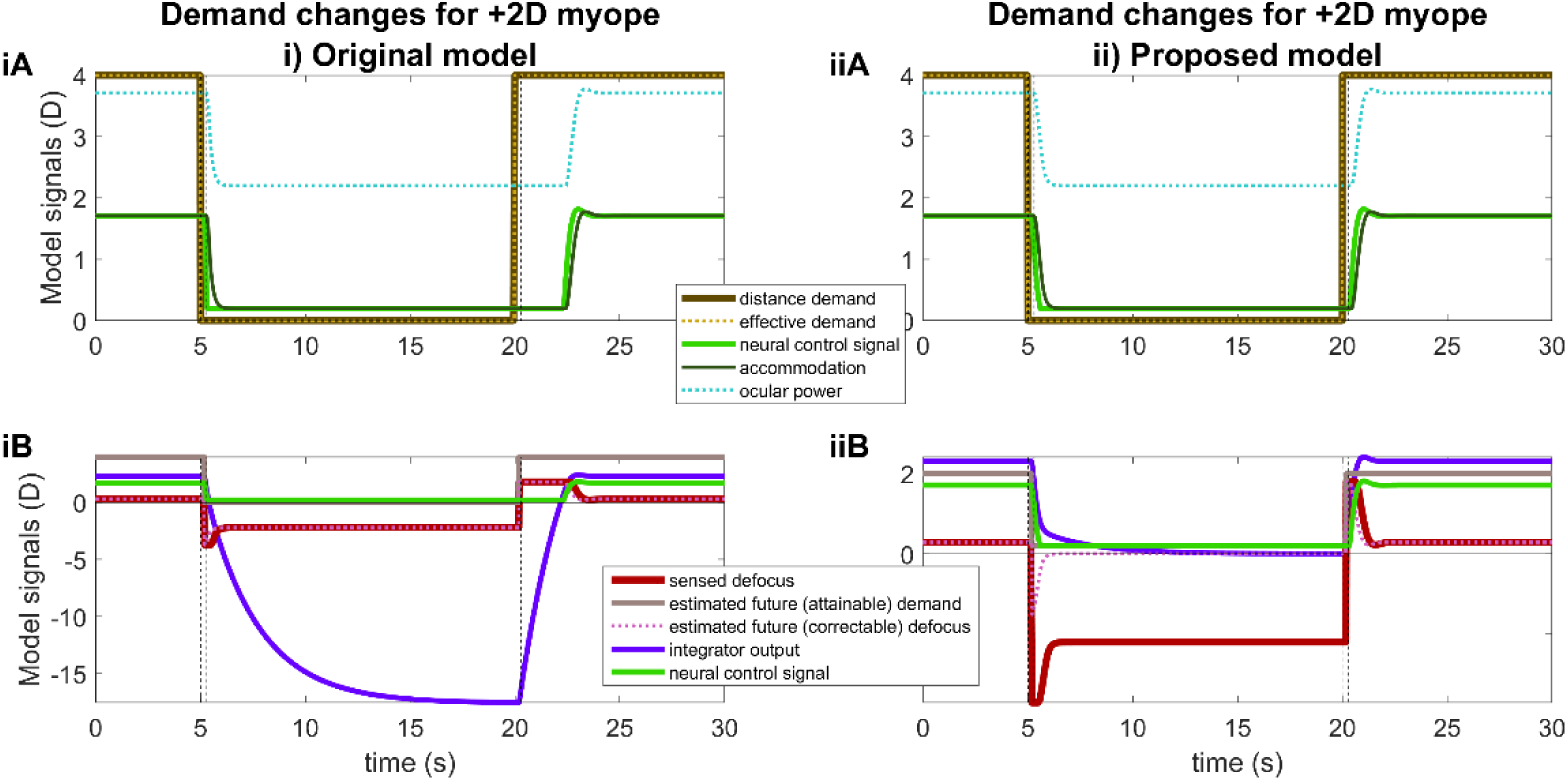
Time-courses of signals for a model myope with +2D refractive error and accommodative range 10D, when stimulus demand steps from 4D to 0D at t=5s and then back from 0D to 4D at t=20s (vertical dashed lines). In this example, the effective demand is equal to the distance demand; we show both for consistency with subsequent figures. The vertical dotted lines mark 300ms after each stimulus change, i.e. the usual sensorimotor latency. The initial relaxation of accommodation occurs with this latency after t=5s, but the subsequent accommodative response is delayed over two seconds beyond the usual latency, because of internal adaptation to the previous tonic defocus error, as shown in B. See Figures 5 and 7 for where the internal signals are taken from in the two models; the same colors are used here as in Figure 7. In the figure legend, “(attainable)” and “(correctable)” apply only to the proposed model (ii, see Figure 7), since the original model (i, see Figure 5) does not make these distinctions. This figure was obtained with Matlab file PlotSignalsForMyope.m

This negative defocus error feeds into the integrator (purple curve). The output of the integrator thus becomes more and more negative, eventually asymptoting to a value around −16D, representing its gain of 8 multiplied by the constant defocus error. The desired command (green curve in Figure 9iB) is equal to the output of the integrator plus a constant bias, here −0.6D (rest focus +1.4D less refractive error +2D). However, the actual motor signal cannot be negative, so remains fixed at 0 (green curve in Figure 9iA). The lens remains fully relaxed, but the refractive error means that the system continues to experience −2D of defocus.

At time t=20s, the stimulus changes back to +4D and after the sensory latency of 200ms, the sensed defocus thus jumps to +2D. The input to the integrator thus becomes positive, and the output of the integrator rises rapidly. However, the motor signal does not become positive until 2s after the increase in demand, and so no change in ocular power or sensed defocus is seen until that point. The model thus predicts that after far viewing, the myope has an enormous, two-second latency before they can refocus on near objects.

In contrast, in the new model (Figure 9ii), the lower bound of the red saturation block shown in Figure 7 means that the estimated future attainable demand is clipped at the minimum neural signal *MinAccomSignal* and is thus +0.2D. Accordingly, while the sensed defocus is −2.2D (red trace in Figure 9iiB), the estimated future correctable defocus is 0D (dashed pink trace in Figure 9iiB: 0.2D attainable demand minus 0.2D accommodation). This means that no signal is fed into the integrator during the period where the myopic eye is experiencing a constant −2D defocus error. The output of the integrator thus falls to 0D during this time (purple trace in Figure 9iiB).

When the stimulus then jumps 2D to +4D at t=20s, the defocus error also jumps 2D, from −2.2D to +1.8D. The estimated future demand relative to refractive error is now +2D, which is within the range of the myopic observer, and so the estimated future attainable demand also becomes +2D. The estimated future correctable defocus therefore transiently becomes close to the actual defocus, +2D. Because the integrator is currently uncharged (value=0), it is able to respond instantly. Accommodation therefore starts increasing as soon as the sensorimotor latency allows, i.e. 300ms after the stimulus change (dark green trace in Figure 9iiA). As accommodation increases, defocus falls and the system enters a new steady-state.

### External lenses

Similar excessive predicted latencies are also seen when refractive error is simulated with external lenses. For example, Figure 10 shows the model predictions for an emmetrope viewing through a +2D “fogging” lens. Initially, the observer is viewing a 0D stimulus and experiencing a steady −2D of defocus blur (red, dotted pink lines in Figure 10B) due to the plus lens and their inability to relax accommodation below 0D. In the original model, this negative defocus error feeds into the integrator, which therefore asymptotes at a large negative value (around −16, representing its gain of 8 multiplied by the constant defocus error of −2D). Subsequently, when the stimulus moves to a distance of +4D, the emmetrope accommodates to +2D, so that the stimulus seen through the +2D lens is in focus. However, due to the previous steady-state defocus and the resulting large negative value of the integrator at the time of the change, there is a delay of >2s before they make this response and thus before sensed defocus decreases.

**Figure 10.**
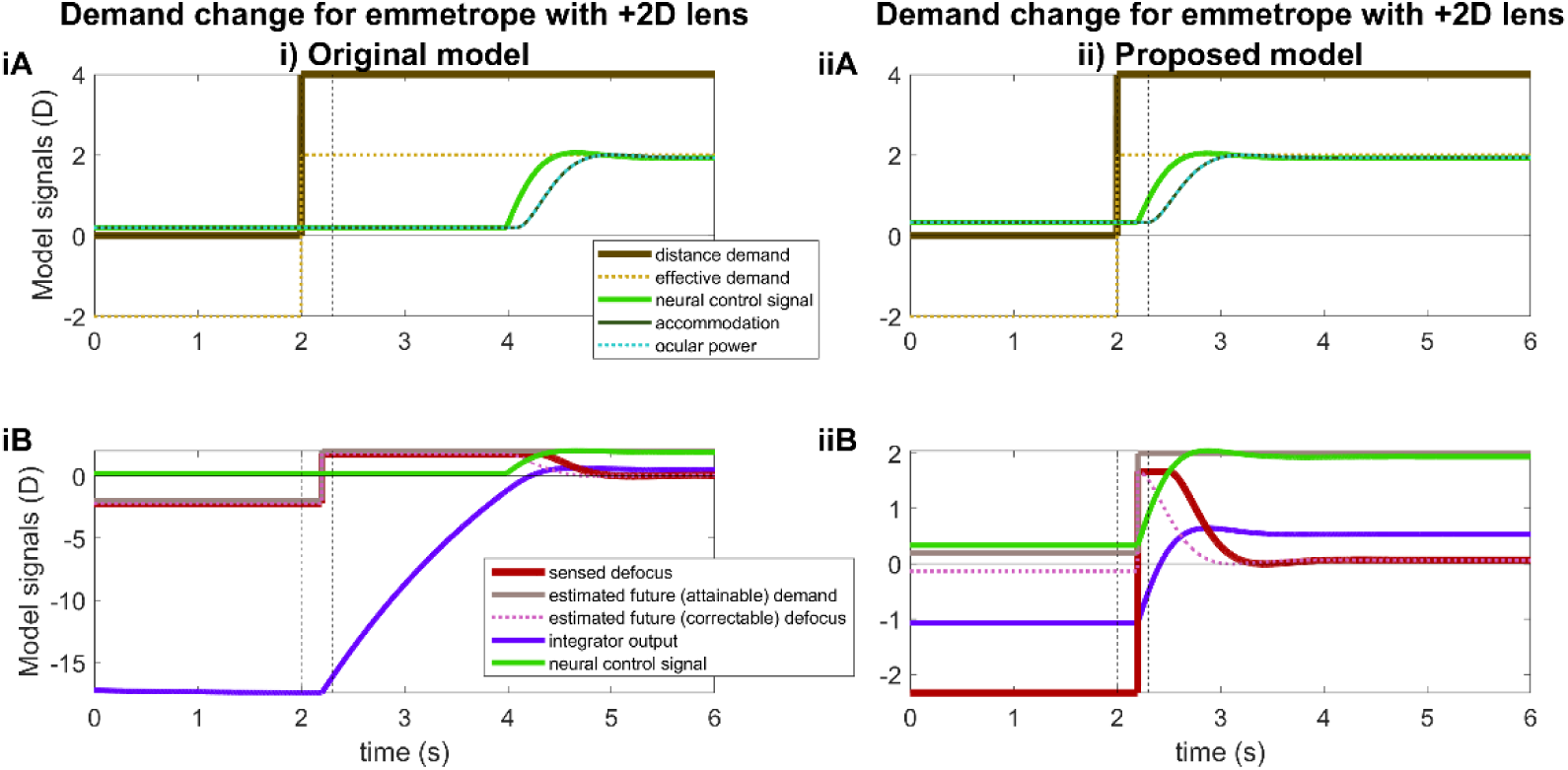
Time-courses of signals for a model emmetrope viewing through a +2D external fogging lens, when stimulus demand steps from 0D to +4D at t=2s (vertical dashed line). The original model (i) predicts a long delay before the emmetrope focuses on the stimulus at 4D, because of adaptation during the preceding period of fogging. Dotted lines mark the time a response would normally begin, i.e. the sensorimotor latency (here 300ms) after the change in demand. Note the different scales of the y axes in iB vs iiB. Other details as for Figure 9. This figure was obtained with Matlab file PlotSignalsForEmmetropePlusLens.m

In the proposed new model, there is again a large optical defocus error exceeding −2D, but the estimated *attainable* demand is just 0.2D, i.e. the minimum neural signal. This ensures that the estimated future correctable defocus remains close to zero (pink dotted line in Figure 10iiB). Thus, the asymptotic output of the integrator is only just below −1D despite the sustained large defocus. Thus, when the stimulus moves closer at t=5s, we see ocular power increasing in response just after the 300ms sensorimotor latency (dashed line at t=5.3s). We do not see the unrealistic 2s latency shown by the original model in Figure 10iA.

Figure 11 shows another example. This is for a model hyperope with a refractive error of −4D and accommodative range of 6D, who can therefore focus on stimuli from −4D to 2D. In this example, the stimulus distance is 4D throughout (heavy brown line in Figure 11A), closer than their maximum ocular power of 2D. Thus when viewed without corrective lenses (t<2s), the demand appears with around +2D of defocus blur even though the hyperope is accommodating as much as they can (accommodation 6D, ocular power 2D). At t=2s, a corrective lens of +4D is applied, fully correcting the hyperopia. The hyperope can now relax accommodation to 4D and focus on the stimulus.

**Figure 11.**
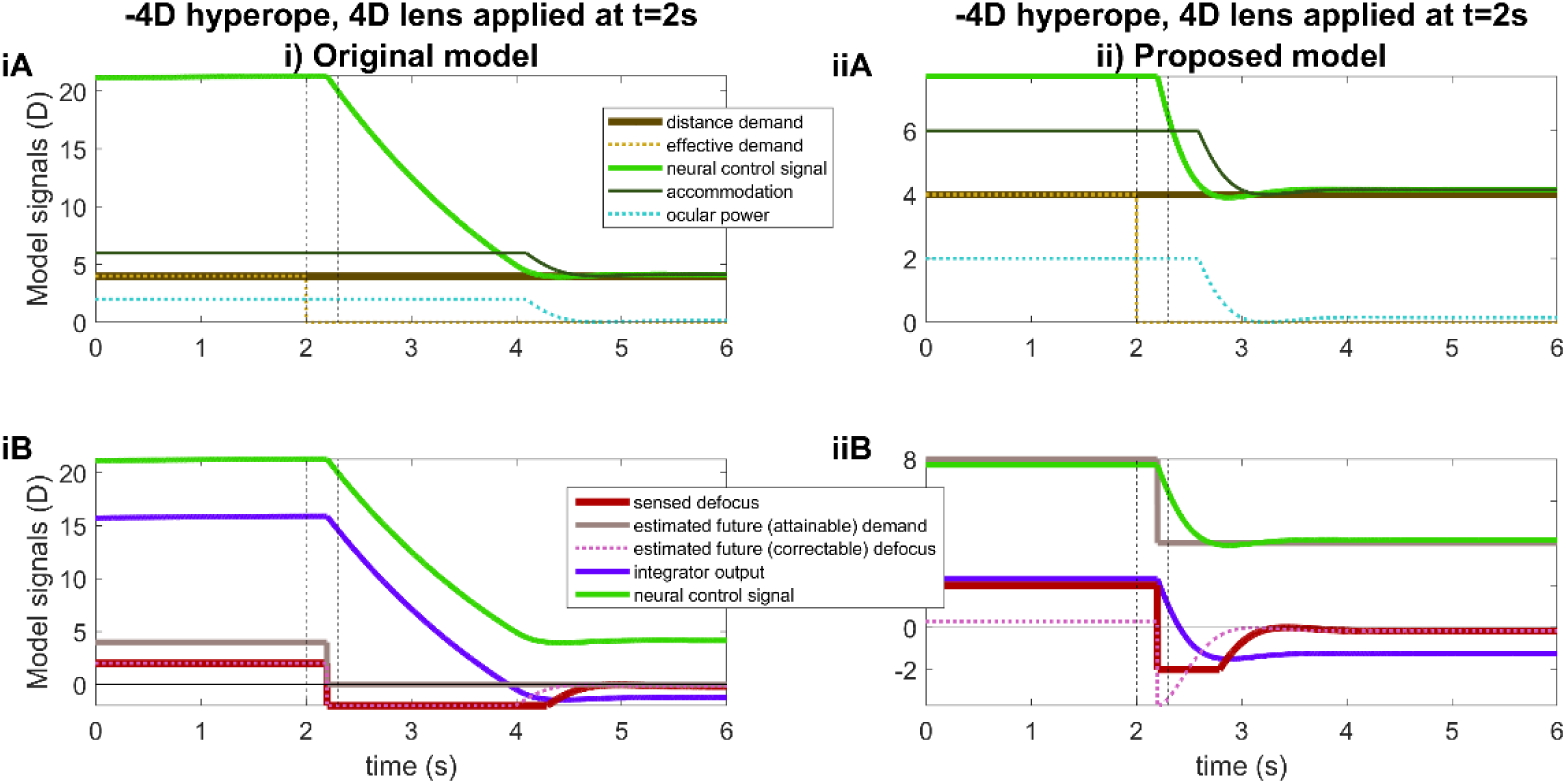
Time-courses of signals for a hyperope with a refractive error of −4D and accommodative range of 6D, before and after a corrective lens is applied at t=2s. The stimulus distance demand is constant at +4D; a corrective lens of +4D is added at t=5s, reducing the effective demand to 0D. Other details as for Figure 9. This figure was obtained with Matlab file PlotSignalsForHyperopePlusLens.m

However, because the integrator is maximally charged following the prolonged defocus (purple trace in Figure 11iB), the original model predicts that no relaxation of accommodation occurs until >2s after the lens is applied (dark green accommodation trace in Figure 11iA). Similarly the sensed defocus jumps from +2D to −2D when the +4D lens is applied, but no reduction in the magnitude of defocus occurs until >2s later (red trace in Figure 11iA). Again, the new model fixes these problems because the integrator was not allowed to charge up to high values. Accordingly, accommodation relaxes shortly after the plus lens is applied, just a little after the 300ms sensorimotor latency (dark green accommodation trace in Figure 11iiA, latency marked with dashed vertical line at 2.3s).

### AC/A ratio

Figure 12 illustrates the problem with the accommodative convergence signal, alluded to in the Introduction. It represents the model’s predictions for an experiment designed to measure the AC/A ratio: the observer fixates monocularly, then the experimenter inserts a minus lens and observes the increase in ocular vergence angle due to the increase in accommodative demand. The stimulus AC/A ratio is the ratio of this change in convergence to the change in stimulus accommodative demand (Alpern et al., 1959; Evans & Pickwell, 2007; Stidwill & Fletcher, 2011). As discussed in the Introduction, in current models the accommodative convergence signal is drawn from the output of the accommodative integrator, multiplied by the AC gain and added onto the output of the vergence integrator (Figure 1). The change in vergence in this experiment is therefore proportional to the change in output of the accommodative integrator produced by the addition of the minus lens.

**Figure 12.**
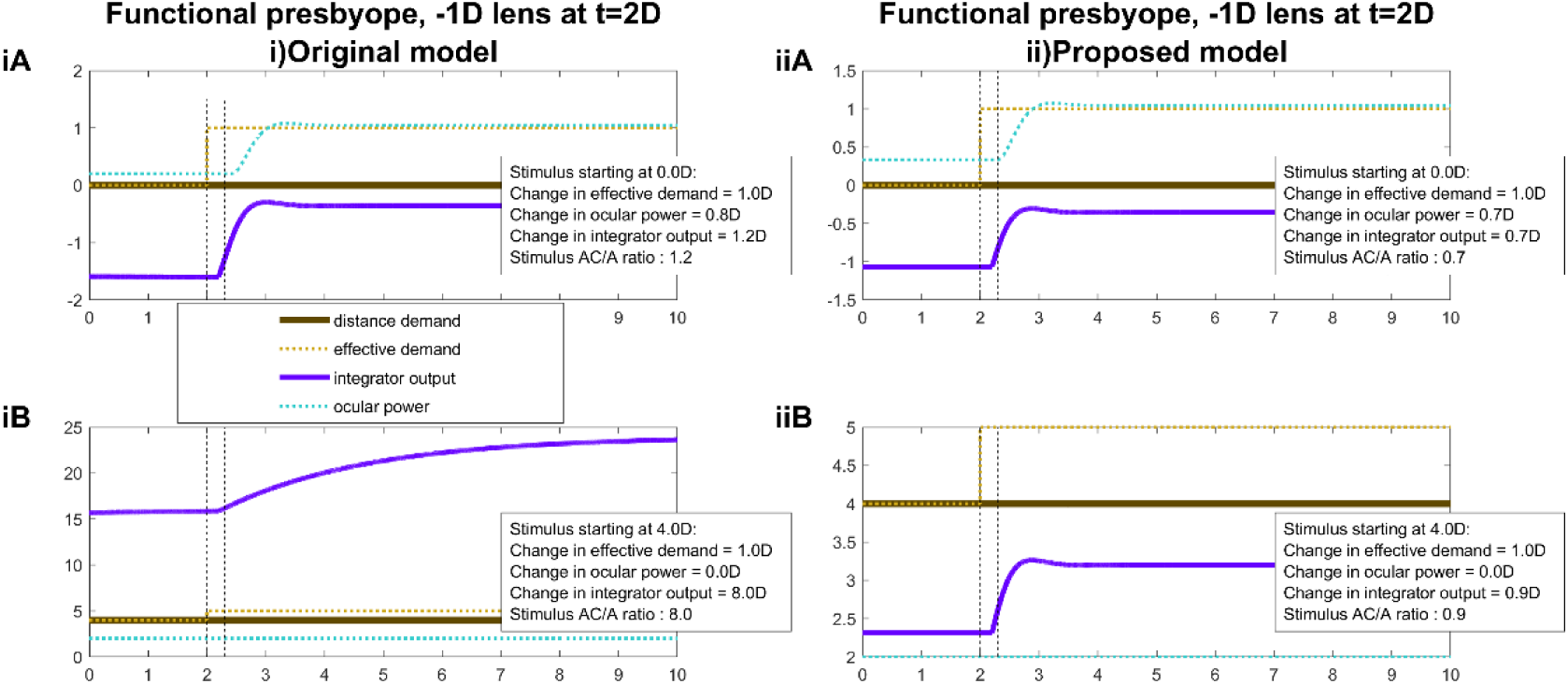
Time-courses of signals for a functional presbyope with a maximum ocular power of 2D, before and after a −1D diverging lens is applied at t=2s, as in an experiment to measure AC/A ratio. The stimulus demand is constant, at (A) infinity, 0D and (B) the observer’s maximum ocular power, 2D. The “stimulus AC/A ratio” is obtained by dividing the total change in integrator output by the change in effective demand. This assumes that the accommodative-convergence signal is the output of the accommodative integrator, as in Figure 1, with unit gain. This figure was obtained with Matlab file PlotACARatio.m

Figure 12iA shows the predictions of the original model when a functional presbyope with maximum ocular power of +2D views a stimulus at 0D, with a −1D lens inserted at t=2s. This all works as expected. When the −1D lens is inserted, the ocular power increases by approximately 1D to compensate, reflecting a similar increase in the integrator output. If the change in convergence reflects the change in integrator output, as in Figure 1, then the stimulus AC/A ratio would be proportional to the change in integrator output. In the figure legend, we have recorded this as “stimulus AC/A ratio”, assuming for simplicity that the constant of proportionality is 1.

However, Figure 12iB shows the problem encountered if the baseline stimulus is closer than the observer’s maximum ocular power of 2D: at 4D, in this example. The observer experiences sustained defocus of +2D, i.e. the difference between the demand of 4D and their maximum accommodation of 2D. This defocus drives the integrator output to around 16D (2D times its gain of 8). Adding the −1D lens increases the defocus to +3D, driving the integrator output up to 24D. This of course has no effect on ocular power, but empirically would produce a further dramatic increase in convergence. Whatever an observer’s AC/A ratio is when the baseline stimulus is within their accommodative range, the current model predicts that it should be much greater for baseline stimuli closer than the near point: multiplied by the gain of the accommodative integrator.

Figure 12ii shows results for the proposed model. Now, the predicted stimulus AC/A ratio increases only marginally when the baseline stimulus is closer than the near point.

The literature does show evidence for a transient increase in AC/A ratio around the maximum ocular power. Figure 13 shows data for two subjects from Alpern et al. (1959). The red disks in Figure 13AB show the ocular accommodation measured during monocular viewing of a stimulus with the effective accommodative demand shown on the x axis. The blue disks show the measured convergence, converted to diopters assuming an interocular distance of 7cm. The blue line in Figure 13CD shows the stimulus gradient AC/A ratio as a function of the initial demand. This is computed by using the smoothed spline lines in Figure 13AB to estimate the increase in ocular convergence when accommodative demand increases by 1D from the initial demand shown in the x axis. The red line shows the same thing for ocular accommodation.

**Figure 13.**
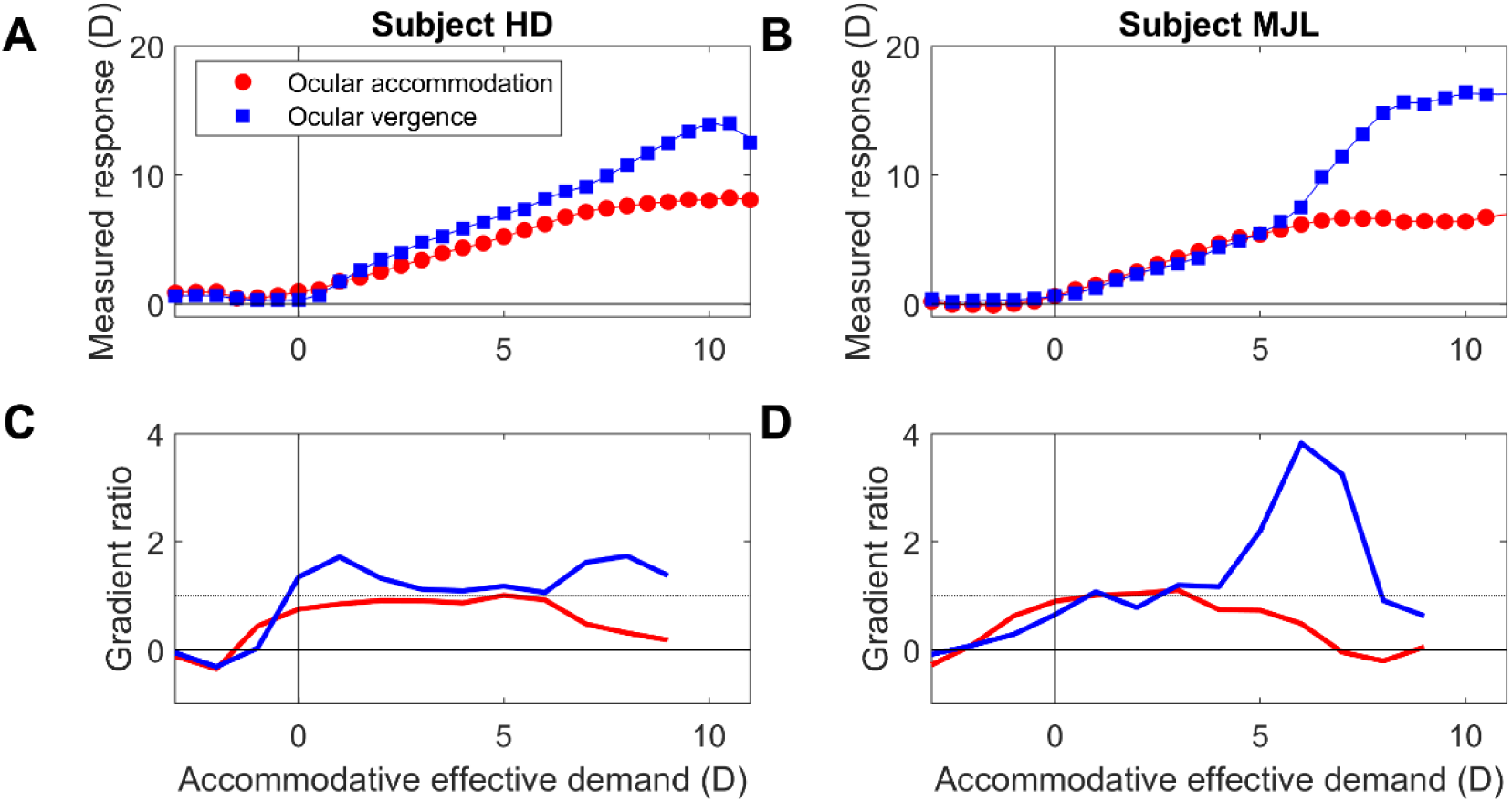
Experimental data digitised from figure 1 of Alpern, et al (1959) for subjects HD (AC) and MJL (BD). AB: Red disks: accommodative response function, i.e. measured ocular accommodation as a function of the effective demand. Blue squares: measured ocular vergence, converted into Diopters to facilitate comparison, assuming an interocular distance of 7cm. This reflects accommodative convergence since the stimulus was monocular. Lines show smoothing splines fitted to data. CD: Red lines: the change in ocular accommodation estimated when accommodative effective demand increases by 1D from the value shown on the x axis, computed using the fitted splines, i.e. an approximation to the derivative of the accommodative response function, designed to be comparable to the stimulus AC/A ratio as computed by the gradient method using a −1D lens. Blue lines: ditto for vergence, i.e. the stimulus AC/A ratio. This figure was obtained with Matlab file Fig_AlpernKincaidLubeck1959.m

Both subjects show AC/A ratios of approximately 1 over most of their accommodative range. However, around the near point, AC/A ratio increases for both subjects. Consider subject MJL (Figure 13BD). This observer has a maximum ocular power of around 6D: beyond that point, his ocular power fails to increase in response to increasing demand. However, his ocular convergence not only continues to increase but from 6-8D it actually rises more steeply for each diopter of accommodative demand, resulting in a nearly fourfold increase in stimulus AC/A ratio. This presumably represents the increase of accommodative effort as the observer strains to focus on the near stimulus, an effect qualitatively similar to that seen in both models (Figure 12). However, this fourfold increase is still lower than plausible for the integrator gain.

Suggestively, however, we do see very large increases in AC/A ratio when accommodation is paralyzed with homatropine to produce a temporary presbyopia. Then, efforts of accommodation are associated with large increases in AC gain (Christoferson & Ogle, 1956). The difference is that here, the range of accommodation is limited pharmacologically, in a way that cannot be known to the neural control system in advance. We present simulations for pharmacological cycloplegia below (Figures 14-17).

**Figure 14.**
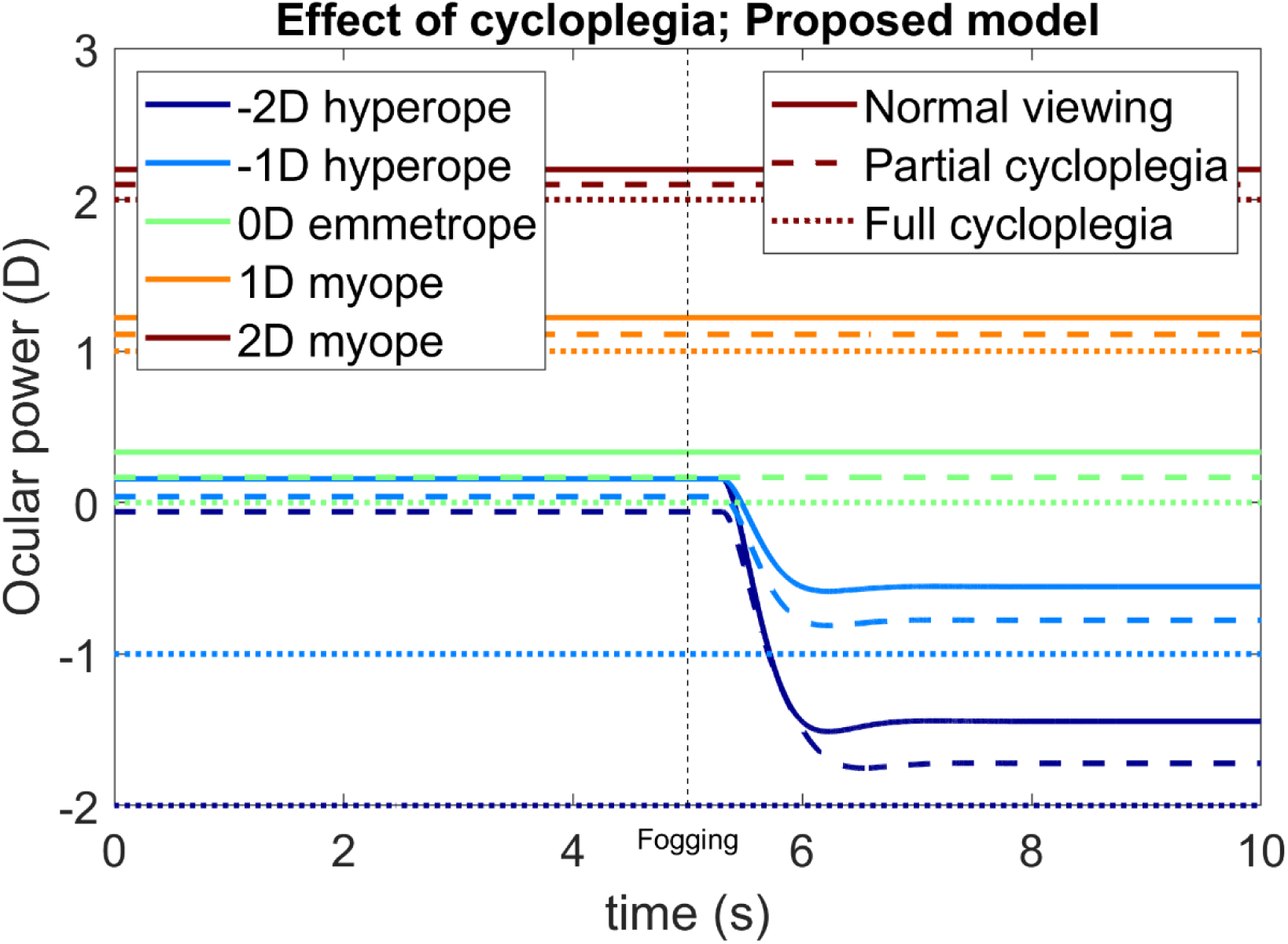
Time-course of accommodation for observers with 5 different refractive errors (color legend) with normal viewing (solid line, model parameter Cycloplegia=0), with partial cycloplegia (dashed line, Cycloplegia=0.5) and with complete cycloplegia (dotted line, Cycloplegia=1). In each case, the observer is viewing a stimulus at infinity (0D) and their accommodative range is 4D. At time t=5s, a +3D fogging lens is added. The model is that shown in Figure 5 (“original model”) with parameters specified in this legend and in Table 1. This figure was obtained with Matlab file PlotOcularPowerCycloplegia.m. Results with the original model are qualitatively similar for this example and are thus not shown in the paper, though can be generated using the Matlab file.

#### Cycloplegia: hyperopic shift

Finally, we consider how to model pharmacological cycloplegia. As noted above, the effect of a cycloplegic drug is modelled as a gain change in the oculomotor nerve signal (triangular gain block in Figures 2 and 3). Importantly, the cycloplegic gain reduction appears in the input to the physical plant, but not in the input to the virtual plant, indicating the brain is unaware of this gain reduction. The cycloplegic effect is described by the model parameter *Cycloplegia*, which ranges from 0 (no cycloplegia, oculomotor nerve signal is unchanged) to 1 (complete cycloplegia, oculomotor nerve signal to the physical plant is abolished). Adding a lower bound to the neural control signal, represented by the model parameter *MinAccomSignal*, enables the model to reproduce the hyperopic shift observed with cycloplegia. The original model behaved sensibly under cycloplegia, so this aspect of the model did not need fixing. We now demonstrate that our proposed new model also gives sensible results.

Figure 14 illustrates the hyperopic shift often seen under partial and complete cycloplegia. This figure shows the time-course of accommodation for five different simulated observers with the refractive errors indicated in the color legend. Results are shown for normal viewing, partial cycloplegia and complete cycloplegia. For each observer, the value of *MinAccomSignal* is 0.2D. The observers are viewing an object at infinity, initially with no external lens and then through a +3D fogging lens applied at t=5s. With complete cycloplegia (dotted line, *Cycloplegia*=1), the value of accommodation is exactly zero and so each observer’s ocular power is simply their refractive error.

In normal viewing with no cycloplegia (solid line, *Cycloplegia*=0), the minimum value of accommodation is *MinAccomSignal*=0.2D. The emmetropic (0D) and myopic (+1D, +2D) observers therefore end up with ocular power 0.2D greater than their respective refractive errors; the addition of the plus lens at t=5s makes no difference as they were already relaxing accommodation as much as possible. The two hyperopic observers, however, initially (during the first 5s) need to accommodate in order to bring the distant stimulus into focus. During this time, their ocular power is a little lower than that of the emmetrope, since it is set by the finite gain of the feedback loop (accommodation is already above the lower bound set by the minimum neural signal). When the +3D lens is applied at t=5s, the two hyperopes relax accommodation and so their ocular power drops. The dashed lines show similar results for partial cycloplegia (*Cycloplegia*=0.5). The key point is that for each simulated observer, the ocular power with cycloplegia is lower than the ocular power measured with distance viewing through plus lenses.

This confirms that placing a lower bound on the signal sent to the oculomotor nerve gives a realistic account of the effect of cycloplegia: cycloplegia produces a hyperopic shift, regardless of the observer’s original refractive error, as observed empirically (Bagheri et al., 2018; Hiraoka et al., 2013; Yoo et al., 2017).

#### Cycloplegia: increase in stimulus AC/A ratio

Figure 15 shows how our model still predicts large stimulus AC/A ratios with cycloplegia, as observed empirically (Christoferson & Ogle, 1956). Figure 15 shows the integrator output and ocular power for an emmetropic observer viewing a stimulus at 0D, before and after the effective demand is increased by the insertion of a −1D lens. Solid lines show the normal responses, and dashed/dotted lines responses when the observer has been partially or completely cyclopleged. In normal viewing (solid lines), ocular power is initially 0.32D, held there by steady-state integrator output of −1.07D combined with the neural bias of 1.4D. Following the insertion of the −1D lens, both integrator output and ocular power increase by around 0.7D. This would imply a stimulus AC/A ratio of around 0.7D (the exact value depending on the AC gain and on the properties of the vergence control system). Because both simulated accommodation and vergence plants are modelled with unity gain, the simulated AC cross-link gain equals the empirically measured response AC/A ratio. The dashed line shows the situation for partial cycloplegia, with the model parameter *Cycloplegia*=0.5. This effectively halves the gain of the system, but of course the virtual plant is unaware of the gain change so the internal estimates of accommodation and demand are double what they should be. The integrator output also doubles as it strives to keep the stimulus in focus despite the lower gain. This halving of the gain thus doubles the AC/A ratio. Finally, dotted lines show the situation for complete cycloplegia, where the feedback loop is severed and ocular power is at 0D throughout. Following the insertion of the −1D lens, the observer experiences sustained defocus of +1D. Critically, this is not predicted by the observer’s accommodative control system, which is expecting the lens to accommodate in response. Thus, the defocus is classed as “correctable” and is fed into the integrator, charging it up to saturation. The stimulus AC/A ratio thus increases by roughly the gain of the integrator relative to normal viewing, just as in the original model.

**Figure 15.**
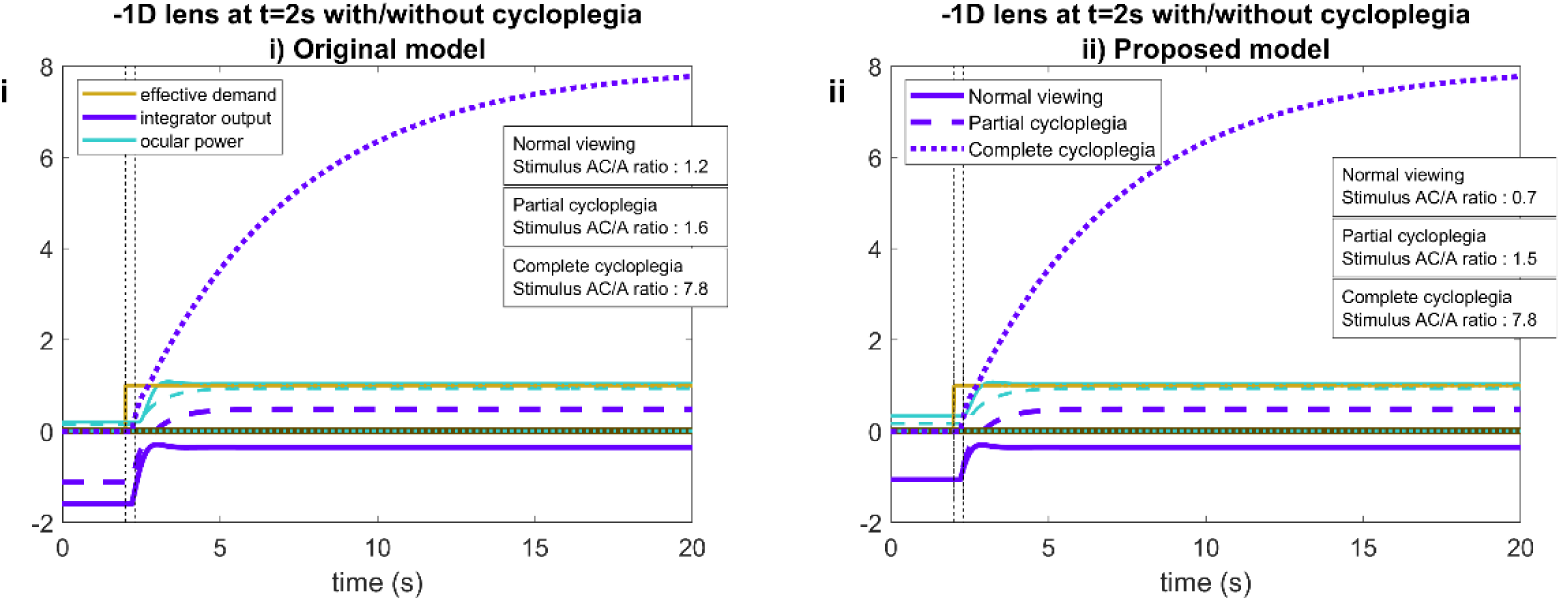
Large AC/A ratio observed with cycloplegia, for both the original and proposed model. Time-courses of signals in (i) the original and (ii) proposed model for an experiment in which an emmetropic observer views a distant stimulus, at 0D, either with normal monocular viewing (solid lines) or with pharmacological cycloplegia (dashed lines). At time t=2s, a −1D lens is inserted, raising the effective demand by 1D. When the observer is cyclopleged, the resulting defocus cannot be corrected. Because this is not predicted by the accommodative control system’s virtual model of the plant, the integrator output becomes very large. This in turn would lead to very large stimulus AC/A ratios. This figure was obtained with PlotACARatioCycloplegia.m

#### Cycloplegia: demand/response curves

We now examine the effect of varying the amount of cycloplegia in more detail. Figure 16A shows the demand/response curve in the presence of different amounts of cycloplegia. As expected, the gradient of the line reduces as cycloplegia increases, becoming flat when cycloplegia is complete. The curves for partial cycloplegia show a distinct “knee”: the gradient becomes much flatter for demands beyond a certain point. This is not observed in current models; there, the initial steep gradient persists throughout. See Appendix for a discussion of why this occurs.

**Figure 16.**
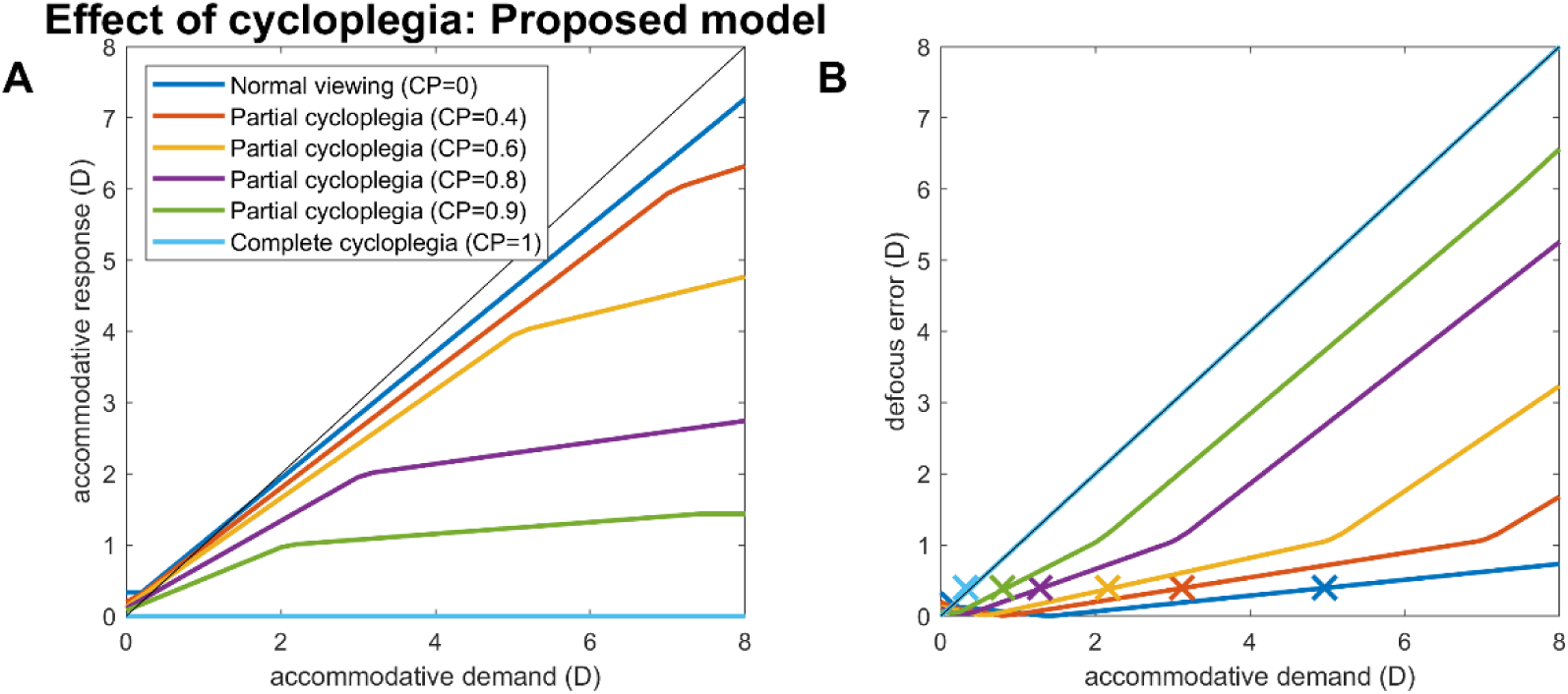
Demand/response curve in the presence of cycloplegia, for the proposed new model. A: steady-state accommodative response for the steady-state demand indicated on the x-axis, for the amount of cycloplegia indicated in the legend. B: magnitude of the steady-state defocus error (i.e. accommodation lag). Crosses x mark the subjective near point assuming a blur threshold of 0.3D, i.e. the highest demand for which steady-state defocus does not exceed 0.3D. Model parameters are as in Table 1 with an accommodative range of 10D and 0D refractive error. This figure was obtained with PlotNearPointCycloplegia.m

Basically, the knee occurs because there are effectively two virtual plants in the internal feedback loop. One has an upper saturation limit of *AccomRange,* AR, and is used to estimate demand and the other has an unlimited upper limit and is used to estimate correctable defocus (see Figures 2 and 7). The knee occurs when the neural control signal exceeds AR. As cycloplegia increases, this point occurs at progressively lower demands, since cycloplegia increases the neural control signal required to elicit a given response. One can show mathematically that the slope is 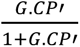 for demands less than 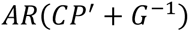 and drops to 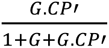 for greater demands, where CP’=1-*Cycloplegia*, AR=*AccomRange* and *G* is the gain of the integrator. In contrast, the original model predicts a slope of 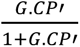 throughout. We are not aware of empirical data which would allow us to test which model gives a better account of human responses.

#### Cycloplegia: reduction in near point

Figure 16B shows how we infer the subjective near point from the simulations shown in Figure 16A. The subjective near point is the distance where observers first notice a blur of the accommodative stimulus, and this is lower than the upper range of accommodation (*AccomRange*, AR). The coloured traces in Figure 16B show the magnitude of the defocus error (the difference between the demand and response). The crosses x show the maximum demand where the defocus remains less than 0.3D, taken to be the threshold for reporting perceptible blur. Figure 17A shows how the inferred subjective near point drops with increasing cycloplegia. Figure 17B shows the stimulus AC/A ratio, prior to the knee, estimated as in Figure 15: that is, while the model observer views a stimulus at infinity, we add in a −1D lens and observe the change in integrator output, for different values of the *Cycloplegia* parameter. Figure 4 of Christoferson & Ogle (1956) similarly plots AC/A ratio and near point for their example observer as a function of time since they administered homatropine. We do not know the mapping from time to the value of the *Cycloplegia* parameter, but we can examine the agreement between model and data by plotting stimulus AC/A ratio (normalized to baseline, since this differs between observers) as a function of near point. This is done in Figure 17C. The solid line shows results for the model simulation; the dots are values for the example observer of Christoferson & Ogle (1956). Qualitatively the agreement is good, with AC/A ratio increasing first slowly, then steeply, as near point reduces. The main discrepancy is that Christoferson & Ogle (1956) found that the maximum AC/A ratio occurred *after* the minimum value of the near point (the highest symbol in Figure 17B is not the leftmost).

**Figure 17.**
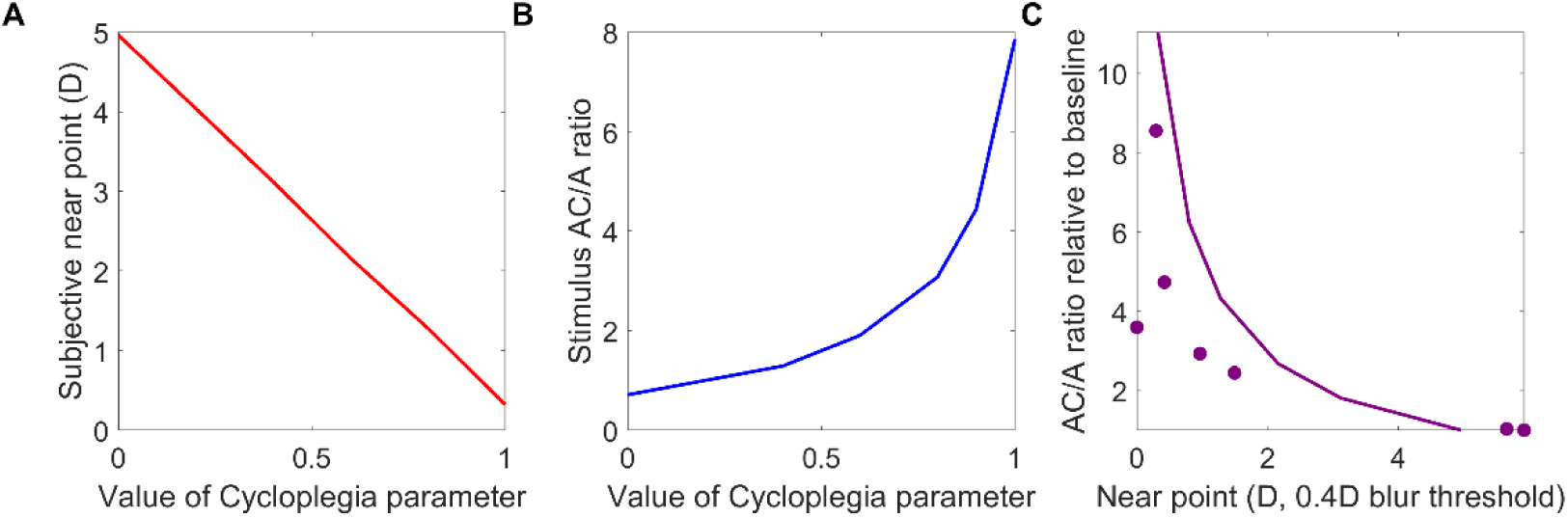
Subjective near point and stimulus AC/A ratios in the presence of cycloplegia. A: Subjective near point (crosses in Figure 16B) as a function of the Cycloplegia model parameter (where 0 indicates normal viewing and 1 complete cycloplegia). B: stimulus AC/A ratio, estimated as the total change in integrator output before vs after a −1D lens is inserted while the model observer views a stimulus at 0D (as in Figure 15), again as a function of Cycloplegia. C: Stimulus AC/A ratio relative to baseline (i.e. divided by the value measured with no cycloplegia) as a function of subjective near-point. Solid line: for the model observer shown in (A); dots: for the representative subject of Christoferson & Ogle (1956); data digitized from their Figure 4. Model parameters are as in Table 1 with an accommodative range of 10D and 0D refractive error. This figure was also obtained with PlotNearPointCycloplegia.m

## Discussion

In this paper, we have shown that conventional ways of incorporating refractive error into models of accommodative control predict unrealistically long response latencies following periods of defocus, whether these are due to refractive error, functional presbyopia, or to fogging with external lenses. This problem with current models has not been pointed out before, presumably because previous work has not happened to examine the response of these models to such situations. We show that this problem can be fixed fairly straightforwardly by adjusting the predictive control such that the signal fed into the integrator is not the predicted future defocus itself, but only that component of predicted future defocus which is correctable. This is just slightly more sophisticated than existing predictive models of accommodative control.

To account for empirical data on accommodation and accommodative convergence, we postulate two different ways of computing “correctable defocus”. For stimuli that are too far, i.e. require ocular power to be reduced below what is possible, we postulate that the brain computes correctable defocus as being the defocus which would result from the actual accommodation if the stimulus were at the furthest attainable distance (but no further). Thus if accommodation is already at the lowest possible value, the correctable defocus is zero. However for stimuli that are too close, i.e. require ocular power to be increased above what is possible, the brain computes correctable defocus as being that which would result from the ideal accommodation (if there were no upper bound to this value) with the stimulus at its actual distance. Thus if the system is already commanding the appropriate accommodation, again the correctable defocus is near-zero (staying at the value needed to obtain the appropriate command), even if the commanded accommodation cannot be obtained. In both cases, therefore, we avoid charging the integrator up with large values of uncorrectable defocus.

We devised these different ways of handling stimuli that are too close vs too far because of the natural asymmetry between these two situations. Due to presbyopia, all humans end up experiencing stimuli that are too close for them to accommodate on. There is value in generating a positive command signal in this situation: both to drive accommodation to the maximum possible, and also to assist convergence. However, an excessive command signal would both produce inappropriate convergence and delay the response to subsequent stimuli. Accordingly, our model generates a positive signal in this situation, but ensures that it stays appropriate to the demand (despite the lack of accommodative response). In contrast, stimuli that are too far to accommodate on should never occur for an emmetropic observer. Commanding a negative accommodation is futile within the accommodation system, and if passed on to the vergence system would produce disruptive divergence. Accordingly, our model avoids generating any signal in this situation. This accords with the clinical experience of author C.S. that myopes do not generally show accommodative divergence when a stimulus already beyond their far point is moved optically still further by the addition of plus lenses.

Note that the model does not require the control system to know its own refractive error. (Refractive error appears in the model but only because we have written the bias signal as “rest focus minus refractive error”. This was for convenience so that, if you know an observer’s measured ocular power at rest focus and their refractive error you can easily infer the value to use for their bias signal. It does not imply that the brain knows its refractive error.) This explains why the new model can solve the latency issue both when defocus is due to the observer’s own refractive error (which could in theory be known to the control system), and also when defocus is due to fogging by an externally applied lens (which could not).

The model does assume that the control system has learnt its own accommodative range. This is not implausible, given that accommodative range changes on a timescale of years, and that predictive control already postulates that the control system has learnt a forward model of its own ocular plant.

We have also extended the model to handle pharmacological cycloplegia, and demonstrate that it gives sensible results for both the subjective near point and the stimulus AC/A ratio in partial as well as complete cycloplegia. This has revealed further interesting nonlinear properties of our model, such as the prediction that with partial cycloplegia, the slope of the demand/response curve will not be reduced uniformly, but will be reduced more strongly for larger demands. This seems reasonably plausible, but has not been tested.

Our postulated modification is fairly simple, plausible and works well in a wide range of clinically relevant situations for accommodation. Whereas the original model showed unrealistically long latencies, we believe our proposed model behaves much more as clinicians would expect. Although we have not simulated full accommodation/vergence models here, we also believe that our modification also makes the output of the accommodative integrator more suitable for use as the accommodative convergence signal in such models.

However, many other approaches are undoubtedly possible. One limitation of our model is that we assume defocus is the sole error function minimised by the system. There could also be other inputs, for example voluntary or involuntary accommodative effort driven by non-optical cues to proximity. For example, the increase in stimulus AC/A ratio reported by Alpern et al (1959) around the end of the accommodative range might reflect an additional voluntary accommodative effort, after the normal involuntary response driven by defocus failed to clear the retinal image.

A similar simplification is that we assume that the predicted future ideal accommodation and attainable demand have no upper bound. More realistically, an upper bound such as the amplitude of accommodation at birth might be used to estimate the ideal attainable bound (Dmax). This could perhaps be quantified from empirical measures of the upper limit to accommodative convergence (ACmax) in absolute presbyopes, assuming that the gain of AC (AC/A ratio) is fixed where Dmax= ACmax/(gain of AC).

Our discovery of this defect in current models has highlighted a need for empirical data on how ocular accommodation behaves during and after exposure to stimuli beyond accommodative range. It is entirely possible that these new data would reveal a need for further modifications to the model proposed here. Our aims in this paper were to (a) draw attention to the gross problem with current models; (b) explain why it occurs; (c) introduce the concepts of correctable defocus, attainable demand and ideal accommodation as offering a potential solution; and finally (d) present a new model that is substantially more accurate than current ones.

We also cannot know whether our model is at all an accurate representation of the underlying neurophysiology (McDougal & Gamlin, 2015). To our knowledge, none of the existing literature can speak to this, since no studies have examined the response to stimuli out of the response range. For example, Zhang et al (1992) identified near response neurons in the midbrain whose firing rate correlates with ocular accommodation for stimuli within the response range of the neuron. In our model, these might correspond to the motor signal itself, or to the estimated future demand, or to the estimated future limited demand, since all these correlate with ocular accommodation for steady-state stimuli. Our model makes different predictions about the behaviour of these three for stimuli out of the response range, but these were not tested. Zhang & Gamlin (1998) found far-response neurons in the posterior interposed nucleus of the cerebellum, whose firing rate increased as accommodation decreased. Microstimulation of this area often elicited decreases in accommodation if accommodation was near, but had no effect if accommodation was already far. Again, this is consistent with our model but hardly diagnostic. For example, these far-response neurons might represent the output of the integrator with a sign inversion; micro stimulation would then request a decrease in accommodation, but the saturation block in the motor signal would explain the lack of response when accommodation is already far. Much more detailed neurophysiology, involving stimuli both within and beyond the range of accommodation, will be needed to elucidate whether this model really captures neural computations. In the meantime, we hope that this descriptive, phenomenological model will prove useful in providing a better account of accommodative responses in a range of clinically relevant situations.

## Acknowledgment

Supported by Magic Leap Inc. by a consultancy contract to Newcastle University for the work of J.C.A.R.

## Appendix: understanding the slope change in the demand/response curve with cycloplegia

For interested readers, we now go through why the change in slope occurs for our new model (Figures 2, 7), and why it does not for the original model (Figure 5). To explain this, it will be helpful to redraw both models in a simplified form suitable for understanding their steady-state response. For this, we can omit all delays such as those due to the sensorimotor latency, since a steady-state signal is unaffected by a delay, and we can replace all transfer functions by their steady-state values. Thus, the physical and virtual plant transfer functions disappear altogether, as does the demand extrapolator, since for all these, in the steady state, the output of the block equals the input to the block and so the block has no effect The integrator, meanwhile, is represented only by its gain G.

In the original model, after these modifications a further simplification becomes possible. Look at the Accommodative Control System in the original model (Figure 5). In the steady state the predicted past and estimated future demands are equal to each other, as are the predicted past and future accommodation signals. Effectively therefore, the signal from the virtual plant is added on and then immediately subtracted. We can thus omit the virtual plant altogether without changing the model’s behavior. This simplified steady-state representation of the model is shown in Figure 18. The predictive nature of the full model (Figure 5) is not relevant in this reduced steady-state version. Effectively, the control system acts directly on the optical defocus (retinal error signal), without reconstructing estimated demand. It is now easy to see that until accommodation reaches its maximum value AR, cycloplegia acts purely to reduce the closed-loop gain of the system. The slope of the demand-response curve is reduced, but there is no “knee”.

**Figure 18.**
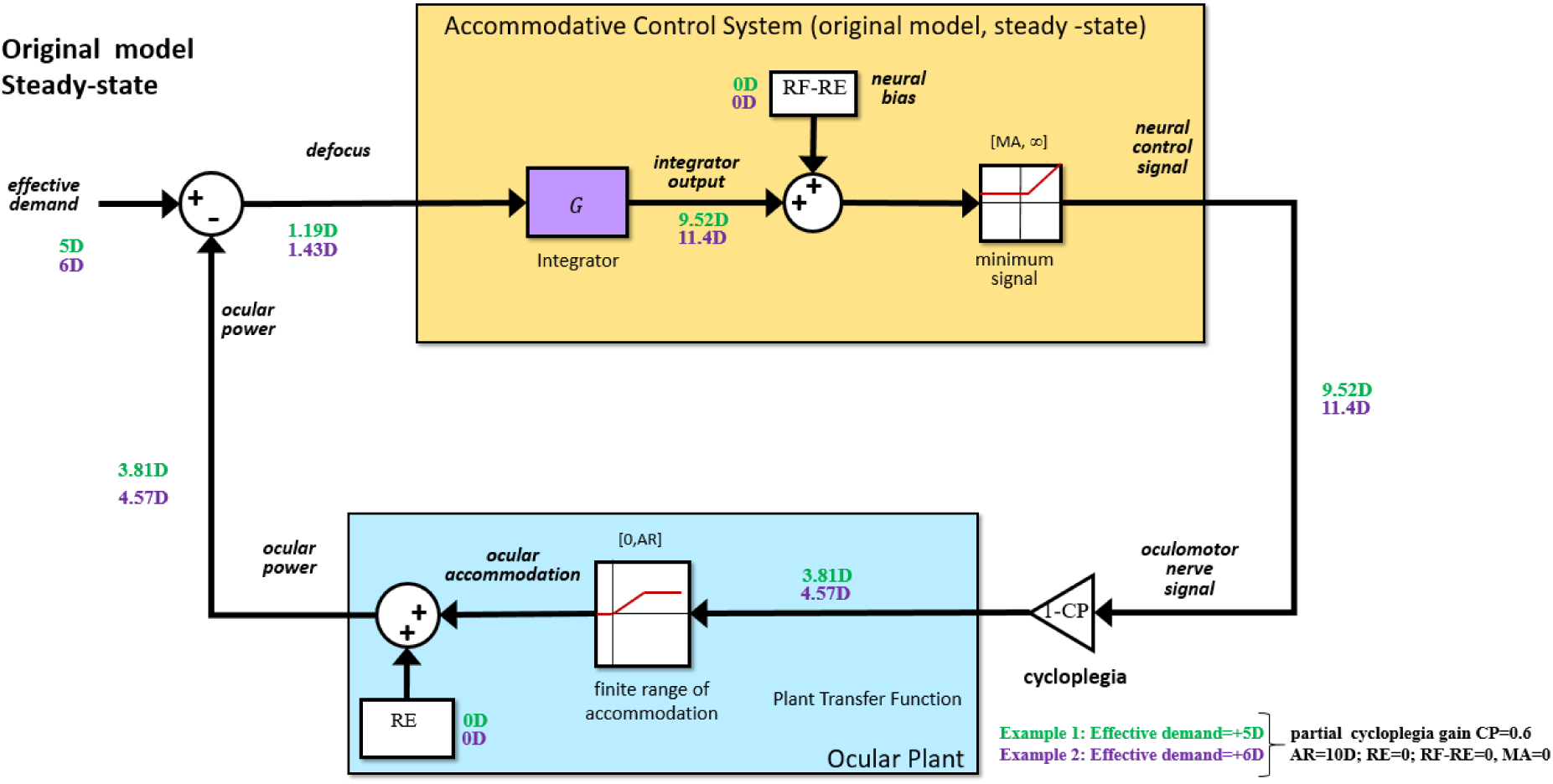
Simplified version of the original model, suitable only for obtaining its steady-state response. Note that in the steady-state, the predictive nature of the full model is irrelevant. Small numbers show signal values for two examples, specified in the bottom right.

The small green and purple numbers work through two simple examples, both with partial cycloplegia (CP=0.6) and for demands of 5D and 6D. The reduced cycloplegic gain reduces the ocular power for a given demand and thus increases defocus. The neural control signal (output of the neural integrator) is thus much larger than it would normally be to achieve the same accommodation: the system is having to “work harder” to achieve even these lower responses. Note that in both examples, the ratio of response to demand is the same, 0.76, reflecting the constant slope of the demand/response curve.

Figure 19 shows a similarly simplified version of our new model, again suitable only for modelling the steady-state. However, note that now the virtual plant cannot be removed. This is because in our new model there are effectively two virtual plants: one incorporating the finite accommodative range of the physical plant, and one not limited in this way. The predicted accommodation signal is the response of the amplitude limited virtual plant, and the predicted ideal accommodation signal is the response of the unlimited virtual plant. Estimated demand is computed by adding predicted accommodation to defocus, and the estimated correctable defocus is computed by subtracting predicted ideal accommodation from estimated attainable demand. Thus, the estimated demand is limited by the amplitude-limited virtual plant and the estimated correctable defocus by the unlimited virtual plant. When the neural control signal (output of the neural integrator that drives the plants) does not exceed the accommodative range of the physical plant, then predicted ideal accommodation and predicted accommodation are equal. When equal, their effects cancel when they are added and subtracted when computing estimated demand and estimated correctable defocus, respectively. Usually, the effective demand (input) is positive relative to refraction, and the estimated attainable demand is equal to the estimated demand. Under these conditions, the estimated correctable defocus equals the optical defocus.

Once the neural control signal exceeds the accommodative range, the predicted accommodation and predicted ideal accommodation will not be equal and thus will not cancel when added to defocus and subtracted from estimated attainable demand, respectively. Additionally, though not relevant here, in general the estimated attainable demand might not be the same as the estimated demand, which could further change results. Consider the example where the neural control signal exceeds the accommodative range of the physical plant. In this example, it is clear from Figure 19 that the amplitude limited predicted accommodation signal will be less than the unlimited predicted ideal accommodation. This means that, for sufficiently large effective demands, the estimated correctable defocus will be less than the optical defocus, whereas for smaller demands they are equal. This effect is what causes the change in slope of the demand/response curve (Figure 16). This illustrates an interesting feature of our proposed new model; its predictive nature remains relevant even in the steady state.

**Figure 19.**
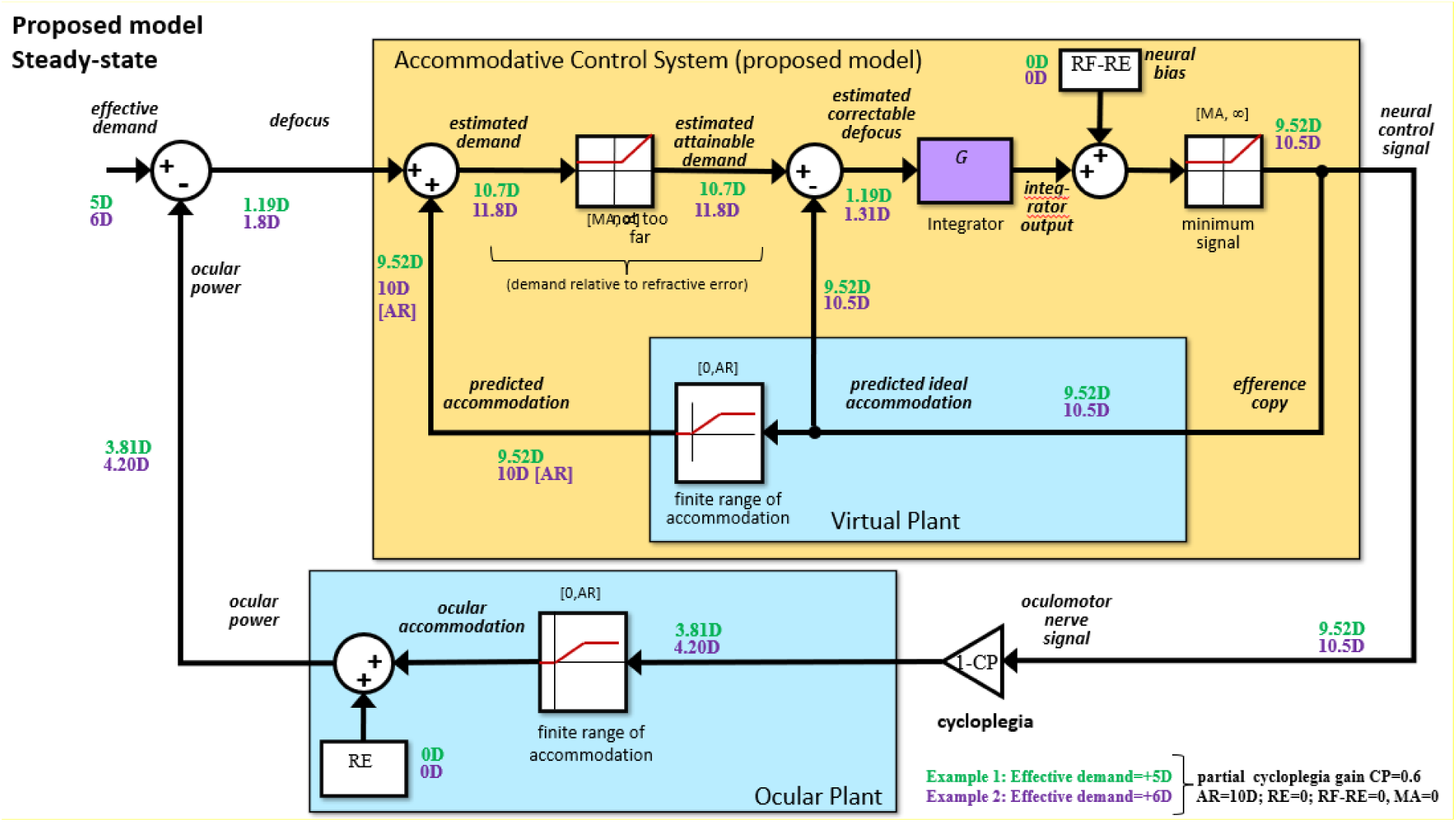
Simplified version of our proposed new model, suitable only for obtaining its steady-state response. The small numbers show the steady-state signal values under partial cycloplegia ( (CP=0.6) for effective demands of 5D (green) and 6D (purple numbers). The accommodative range is AR=10D and the gain G=8. As before in these worked examples, the neural bias is set to zero for simplicity (RF-RE=0). With the specified values, the “knee” is 5.25D, so the two example demands are on either side of this value.

To go through this in detail, the small numbers in Figure 19 show steady-state values for two different demands, 5D and 6D, with partial cycloplegia. The accommodative range is AccomRange=10D and Cycloplegia=0.6, so the change in slope occurs at 5.25D. The 5D demand (green numbers) is below this point, so our new model responds exactly the same as the original model (compare green numbers in Figure 18 vs 19). However, now consider the situation for the 6D demand (purple numbers). Here, the accommodation under partial cycloplegia is 4.2D. Because of the cycloplegic gain reduction, to achieve this value, accommodation requires a neural control signal of 10.5D, beyond the range of accommodation (10D). The brain thus estimates the attainable demand as 11.8D (10D predicted accommodation, estimated by the amplitude-limited virtual plant, plus the 1.8D defocus). This is actually slightly higher than the 11.4D estimated for the 6D demand the original model (Figure 18), due to the higher optical defocus in our model. However, to compute the estimated correctable defocus, our model then subtracts the predicted ideal accommodation of 10.5D – effectively the signal from a second virtual plant with unlimited amplitude. Because the 10.5D signal subtracted from this unlimited virtual plant is larger than the 10D signal originally added on from the limited virtual plant, the estimated correctable defocus signal, 1.3D, ends up both being lower than the actual defocus, 1.8D, and lower than the defocus fed into the integrator in the original model, 1.43D (Figure 18). Thus, the effective gain of the system is now reduced even more than implied by the cycloplegic gain reduction. This nonlinear effect accounts for the change in slope of the demand/response curve (Figure 16): from 0.76D change in response for each 1D change in demand for stimuli further than 5.25D, to 0.26D change in response per 1D change in demand for nearer stimuli.

